# Metabolic and mitochondria alterations induced by SARS-CoV-2 accessory proteins ORF3a, ORF9b, ORF9c and ORF10

**DOI:** 10.1101/2023.09.26.559506

**Authors:** Blanca D. López-Ayllón, Silvia Marín, Marco Fariñas Fernández, Tránsito García-García, Raúl Fernández-Rodríguez, Ana de Lucas-Rius, Natalia Redondo, Laura Mendoza-García, Carles Foguet, Juozas Grigas, Alba Calvet, José Manuel Villalba, María Josefa Rodríguez Gómez, Diego Megías, Biagio Mandracchia, Daniel Luque, Juan José Lozano, Cristina Calvo, Timothy M. Thomson, Juan J. Garrido, Marta Cascante, María Montoya

## Abstract

Antiviral signaling, immune response and cell metabolism in human body are dysregulated by SARS-CoV-2, the causative agent of the COVID-19. Here, we show that SARS-CoV-2 accessory proteins ORF3a, ORF9b, ORF9c and ORF10 induce a significant mitochondrial and metabolic reprogramming in A549 lung epithelial cells. While all four ORFs caused mitochondrial fragmentation and altered mitochondrial function, only ORF3a and ORF9c induced a marked structural alteration in mitochondrial cristae. ORF9b, ORF9c and ORF10 induced largely overlapping transcriptomes. In contrast, ORF3a induced a distinct transcriptome, including the downregulation of numerous genes for proteins with critical mitochondrial functions and morphology. Genome-Scale Metabolic Models predicted common and private metabolic flux reprogramming, notably a depressed amino acid metabolism, and an enhanced metabolism of specific lipids distinctly induced by ORF3a. These findings reveal metabolic dependencies and vulnerabilities prompted by SARS-CoV-2 accessory proteins that may be exploited to identify new targets for intervention.

**One-Sentence Summary:** Mitochondria and metabolic alterations induced by SARS-CoV-2 accessory proteins ORF3a, ORF9b, ORF9c, ORF10 in pulmonary cells unravel new targets of intervention.

## INTRODUCTION

Severe acute respiratory syndrome coronavirus 2 (SARS-CoV-2) is responsible for over 750 million cases of coronavirus disease 2019 (COVID-19) and over 7 million deaths since its emergence in 2019/2020^1^. Upon infecting susceptible cells, SARS-CoV- 2 subverts and disables innate immune responses and, at the same time, hijacks metabolic pathways to favor viral replication and production at the expense of the infected cell. As such, metabolic reprogramming is the most consistent molecular change in SARS-CoV-2 infection^2^. Affected metabolic pathways including lipid metabolism, glucose metabolism, and oxidative phosphorylation have been observed in human patient samples, lung airway and alveolar organoids and metabolic models^2, 3, 4, 5^. Although a complete mechanism explaining the development of all the aforementioned metabolic alterations in COVID-19 is yet to be proposed, viral-host protein interactions and the specific subcellular localization of various SARS-CoV-2 components help to identify potential sites and pathways leading to COVID-19 pathogenicity.

In particular, mitochondrion, an organelle which plays an important role in maintenance of multiple metabolic functions, such as energy metabolism and reactive oxygen species (ROS) production, has been recognized as a relevant organelle for the life cycle and cytopathic effects of SARS-CoV-2, as indicated by an enrichment of viral dsRNA signal in mitochondria, coupled with mitochondrial lesions caused by SARS-CoV-2 infection ^6, 7^. Furthermore, changes in mitochondrial shape and structure, cristae reorganization and membrane potential disruption have been observed in SARS-CoV-2- infected cells ^6^, as well as inhibition of mitophagy with consequent accumulation of damaged mitochondria and augmented stress signaling by inhibiting LC3 binding ^7^. By contrast, other study suggests that ORF10 interacts with mitophagy receptor Nip3-like protein X (NIX) and LC3, in turn triggering mitophagy which hinders membrane-anchored anti-viral signaling protein (MAVS)-mediated antiviral signalling ^17^. In addition to their metabolic and bioenergetic functions, mitochondria play an integral role in host innate immune responses and interferon signaling. In particular, recognition of viral RNA by cytoplasmic sensors is followed by interaction with, and activation of MAVS ^8,9^.

In addition to 4 main structural and 16 nonstructural proteins, the SARS-CoV-2 genome encodes 11 accessory proteins^10, 11^. Although accessory proteins were initially thought as minor contributors to the survival and pathogenesis of the virus, multiple studies point out their roles in immune response evasion strategies, interactions with metabolic pathways and induction of host cell apoptosis^10, 12, 13^. ORF3a, a membrane-associated protein and the largest SARS-CoV-2 accessory protein, has been reported to induce apoptosis in cell culture models^14^ and to modulate host innate immune responses^15^, notably a suppression of IFN-I signaling via impeding signal transducer and activator of transcription 1 (STAT1) activation and nuclear translocation, resulting in increased suppressor of cytokine signaling 1 (SOCS1) levels^15^; however, its role as a viroporin remains controversial^16^. Other SARS-CoV-2 accessory proteins, such as ORF9b and ORF10, also suppress IFN-I responses^12, 17^. ORF9b inhibits the binding of translocase of the outer membrane 70 (TOM70) to heat shock protein 90 (Hsp90), which is crucial for Hsp90/TANK-binding kinase 1 (TBK1)/interferon regulatory factor 3 (IRF3) complex recruitment on mitochondria, resulting in suppression of antiviral gene transcription^12^. ORF9c interacts with Sigma receptors, implicated in lipid remodeling and stress responses in the endoplasmic reticulum^18^. Also, ORF9b and ORF9c have been involved in inflammatory and fibrotic processes via IL-11^19^.

Prior evidence for SARS-CoV-2-induced metabolic changes and the putative functions ascribed to accessory proteins ORF3a, ORF9b, ORF9c and ORF10 prompted us to investigate the potential for metabolic reprogramming by these ORFs. To this end, A549 human lung carcinoma cells were individually transduced with ORF3a, ORF9b, ORF9c and ORF10, analyzed for mitochondrial morphology, mass and function, and transcriptomic and metabolomic data generated in order to build genome-scale metabolic models (GSSM). We have found that SARS-CoV-2 accessory proteins ORF3a, ORF9b, ORF9c and ORF10 induced significant mitochondrial alterations and metabolic reprogramming. Further, GSMMs predicted common and private metabolic flux reprogramming, notably depressed amino acid metabolism and enhanced lipid metabolism distinctly induced by ORF3a. These findings reveal metabolic dependencies and vulnerabilities induced by specific SARS-CoV-2 accessory proteins, that might be exploited to identify new targets of intervention.

## RESULTS

### ORF3a and ORF9c induce significant structural alterations in mitochondria

SARS-CoV-2 accessory proteins, ORF3a, ORF9b and ORF9c, have been shown to associate with mitochondria^20, 21^. In order to further understand the consequences of the expression of these proteins on mitochondrial function and cellular metabolism, these proteins were individually transduced into A549 lung carcinoma cells (ORF-A549 cells) and drug-selected for stable constitutive expression of the viral genes (Fig. S1). As controls, non-transduced A549 cells and cells transduced with GFP only-expressing lentiviruses were used. Both controls yielded equivalent results in all assays used throughout this study (data not shown).

Ultrastructural analysis by transmission electron microscopy showed a shorter mitochondria length in all ORF-A549 cells, except in ORF9b-A549, as compared to control cells (Fig. 2A, 2B, Fig. S2 and Fig S3). Moreover, a lower number of ridges were observed in most ORF-A549 cells, with a more diffuse appearance (Fig. 2C, Fig. S2 and Fig S4), conforming less organized cristae with a loss of parallelism and more hollow mitochondria than in control cells (Fig. 2A and Fig. S2). Parallelism in the cristae was defined as cristae aligned in the same orientation. All ORF-A549 cells also displayed filamentous bundles frequently located around the nucleus, which were not usually observed in control cells.

**Figure 1.**
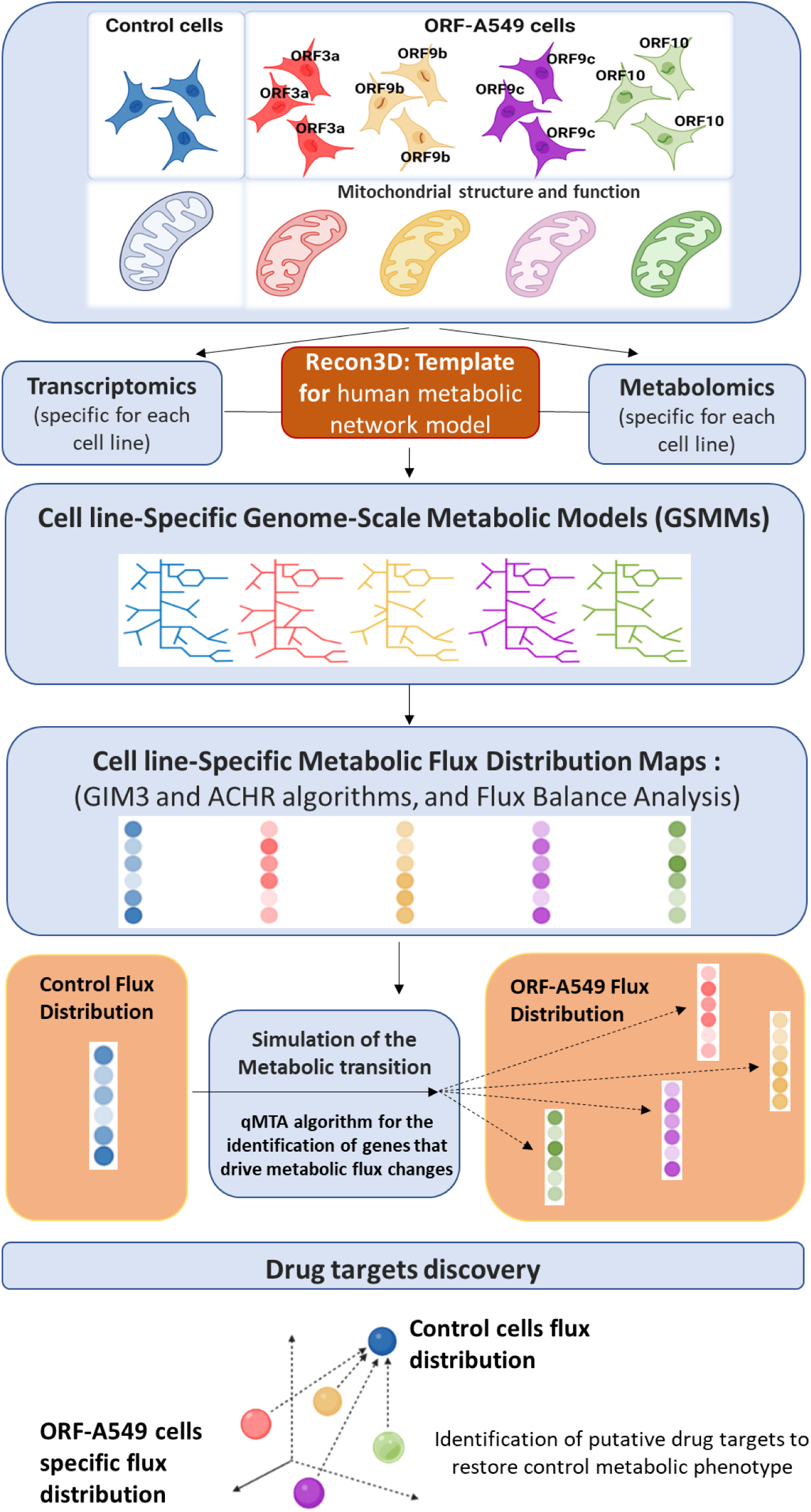
Experimental workflow. Image generated in Biorender.

**Figure 2.**
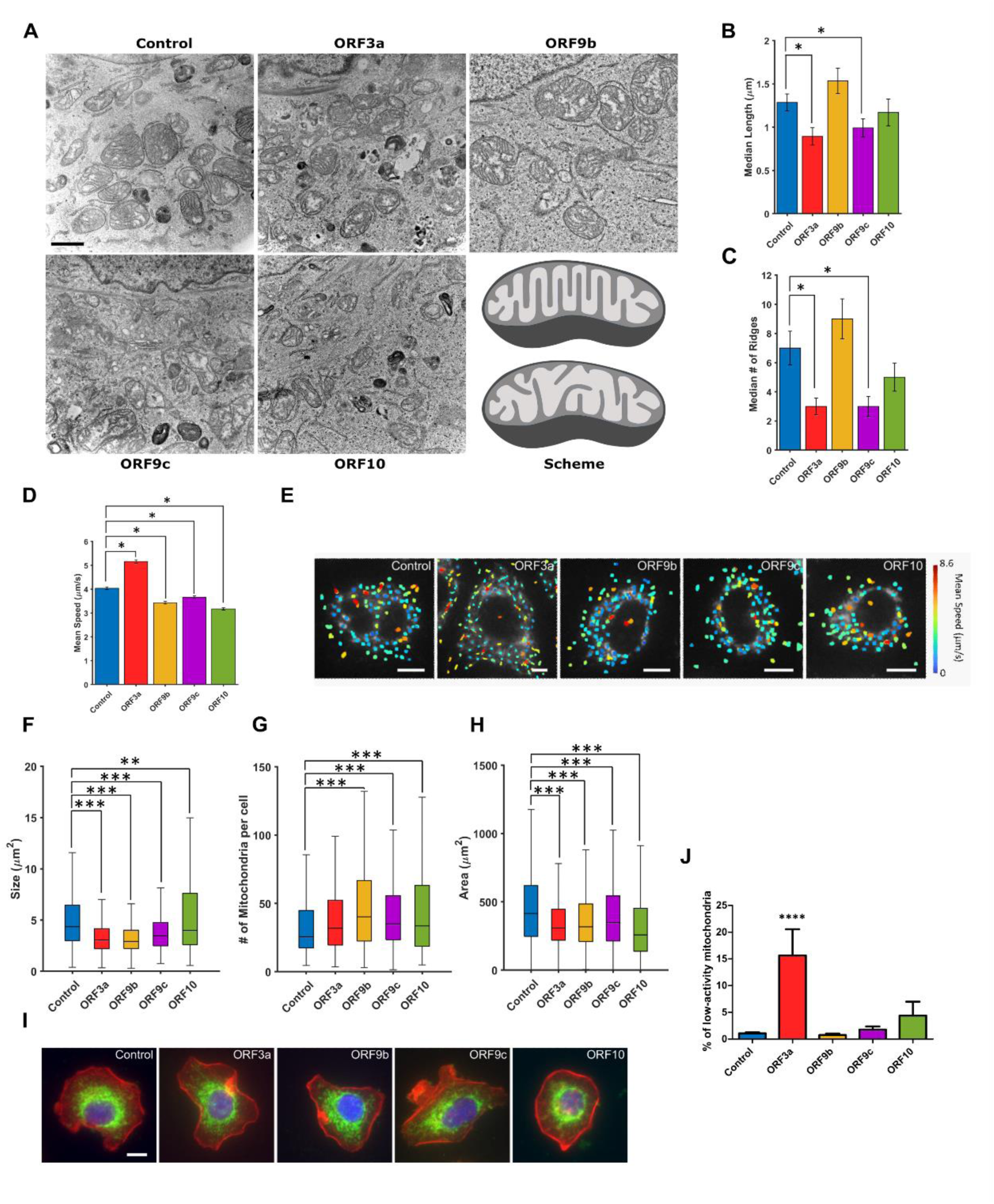
Mitochondrial morphology and diffusion in ORF-A549 cells. **(A)** Analysis by transmission electron microscopy of mitochondrial appearance in ORF-A549 cells ultrathin sections. Scale bar indicates 1µm. Last panel shows a schematic representation of mitochondria with organized and parallel mitochondrial ridges (top), and mitochondria with disorganized, non-parallel mitochondrial ridges and hollowed out areas (bottom). Median of mitochondria longest diameter **(B)** and median number of ridges per mitochondrion **(C)** evaluated from TEM micrographs of each cell line. **(D)** Mean speed of the tracked mitochondria measured for each cell line. **(E)** Representative results of the tracking performed on fluorescently labeled mitochondria. Taking the irregular shape of mitochondria into account, a mask-based segmentation to separate the different organelles was used. Track colors represent the average speed measured in between two frames. Scale bars: 10 µm. **(F-H)** Box and whisker plots showing the distributions of mitochondrial size (mitochondria fragmentation) **(F)**, number per cell **(G)**, and area covered by mitochondria (mitochondrial mass) **(H)**. **(I)** Representative images of the datasets used to generate the distributions in **(F-H)**. Mitochondria are marked in green (TOM20), nuclei in blue (DAPI), and cell membrane in red (Phalloidin). Scale bar: 10 µm. **(J)** Quantification of the percentage of low-activity mitochondria in ORF-A549 cells stained with MitoView 633. Data are represented as mean ± SD (n=4). Statistical significance is as follows: *p < 0.05, **p < 0.01, ***p<0.001, ****p < 0,0001.

As mitochondrial morphology, function and motility are coupled^22, 23^ a study of mitochondrial motility was next performed by live-cell imaging (Fig. 2E). Analysis of mitochondria motion showed significant differences among the different ORF-expressing cells (Fig. 2D). Based on their motility, two subpopulations of motile mitochondria were observed in all cell lines, namely low-and high-speed mitochondria, in addition to stationary mitochondria. In ORF3a-A549 cells, the number of mitochondria traveling at high speed was significantly higher than in control cells (Fig. S5). In the rest of ORF-A549 cells, the number of mitochondria traveling at low speeds was significantly higher than in control cells (Fig. S5).

It is worth noting that the size of mitochondria was smaller in all ORF-transduced cells compared to control cells, being particularly evident in ORF3a-A549 and ORF9b-A549 cells (Fig. 2F and 2I), indicative of augmented mitochondrial fission. Moreover, ORF-A549 cells showed an increased number of mitochondria compared to control cells, although in the case of ORF3a-A549 it was not statistically significant (Fig. 2G and 2I), and total cell area covered by mitochondria (mitochondrial mass) was smaller in all four ORF-A549 cells versus control cells (Fig. 2H, 2I). Given the effect of ORFs on mitochondria morphology, mitochondrial membrane potential (Δψ_m_) was analyzed by flow cytometry using MitoView 633 (Fig. 2J and Fig. S6). Two populations were defined according to their mitochondrial membrane potential: high-activity and low-activity mitochondria. Indeed, ORF3a-A549 cells exhibited a significantly higher number of low-activity mitochondria in their cytoplasm. ORF10-A549 cells also showed an increase in this mitochondrial population, although without reaching statistical significance.

In summary, ORF3a favors mitochondrial fission and altered cristae architecture, coupled to diminished Δψ_m_ and enhanced motility in a significant population of mitochondria. On the other hand, ORF9c, while causing mitochondrial fission and ultrastructural alterations similar to those induced by ORF3a, did not cause loss of Δψ_m_ or enhanced mitochondrial motility. ORF9b and ORF10 also induced mitochondrial fragmentation and structural alterations albeit less significantly than ORF3a or ORF9c.

### ORF3a induces a highly divergent transcriptome in A549 cells

In order to shed light on the biochemical pathways and biological processes modulated by ORF3a, ORF9b, ORF9c and ORF10, a comparative transcriptomic analysis of ORF-A549 cells was performed. Unsupervised hierarchical clustering of all detectable transcripts showed that ORF3a-A549 cells displayed a clearly distinct transcriptome from control cells and from the rest of ORF-A549 cells (Fig. S7 and S8A). Transcriptomes of cells expressing ORF9b, ORF9c or ORF10 were less distinct, overall, from control cells or from each other (Fig. S7and S8A), although one-to-one comparisons of each ORF-A549 to control cell transcriptomes yielded a number of significant differentially expressed genes (DEGs) (Supplementary Table 1). Several transcripts significantly upregulated in ORF3a-A549 cells, but not in cells expressing the other three ORFs, were linked to innate immune functions, including interferon-regulated genes such as IFI44L pattern-recognition receptors (PRRs) such as TLR5, or the complement and coagulation cascades, including VWA2, F5 or F13B (Supplementary Table 1). Likewise, a number of transcripts coding for proteins with metabolic functions were significantly overexpressed in these cells, including transporters (e. g., SLC9A9, SLC16A7, SLC16A9 or SLC25A27) and enzymatic metabolic regulators (e. g., ADH4 or ACSL5) (Supplementary Table 1). On the other hand, we observed a significant overlap of up-or down-regulated DEGs among cells expressing ORF9b, ORF9c and ORF10 cells, with over 30% transcripts shared by each with the two other gene sets (Supplementary Table 1). Outstanding among the downregulated transcripts shared among cells expressing ORF9b, ORF9c and ORF10 cells, we found Stimulator of Interferon Response cGAMP Interactor 1 (STING1), accompanied with IFN signaling pathway genes such as IRF6, IFITM1 or IFITM3 in ORF9c-A549 cells or IFI44 or IFITM1 in ORF10-A549 cells, consistent with known functions exerted by these viral proteins as IFN-I and III signaling antagonists^24, 25^.,

We next resorted to gene-set expression analysis (GSEA)^26^, in order to gain a broader perspective of biochemical pathways and cellular processes perturbed by the expression of these viral accessory proteins (Fig. S8B-E). Focusing on metabolic pathways revealed that ORF3a, but not the other ORFs, induced significant changes in specific metabolic pathways, including enrichment of the biliary acid pathway and underrepresentation of the mTORC1, hypoxia, glycolysis, oxidative phosphorylation (OXPHOS) or mitochondrial reactive oxygen species (ROS) pathways (Fig. S8B and 3). Interestingly, many of the genes in the biliary acid pathway upregulated by ORF3a code for peroxisome biogenesis proteins, such as PEX1, PEX6, PEX7, PEX11A, PEX11G, PEX12, PEX13, PEX16 and PEX17 (Fig. 3). This suggests a coordinated upregulation of these peroxisomal biogenesis genes and is in line with the increased abundance of peroxisomes observed upon infection of lung epithelial cells with SARS-CoV-2 ^27, 28^. The inference of enhanced peroxisomal abundance and/or function induced by ORF3a is further supported by the additional upregulation of numerous genes for peroxisomal enzymes (PIPOX, NUDT12, HAO1, ISOC1, PRDX5, PHYH, GSTK1, AMACR, HSD17B4, PAOX, PNPLA8, SCP2, EPHX2, CROT) and transporters (ABCD1, ABCD2, ABCG4) involved in sterol and bile acid synthesis. (Fig. 3).

**Figure 3.**
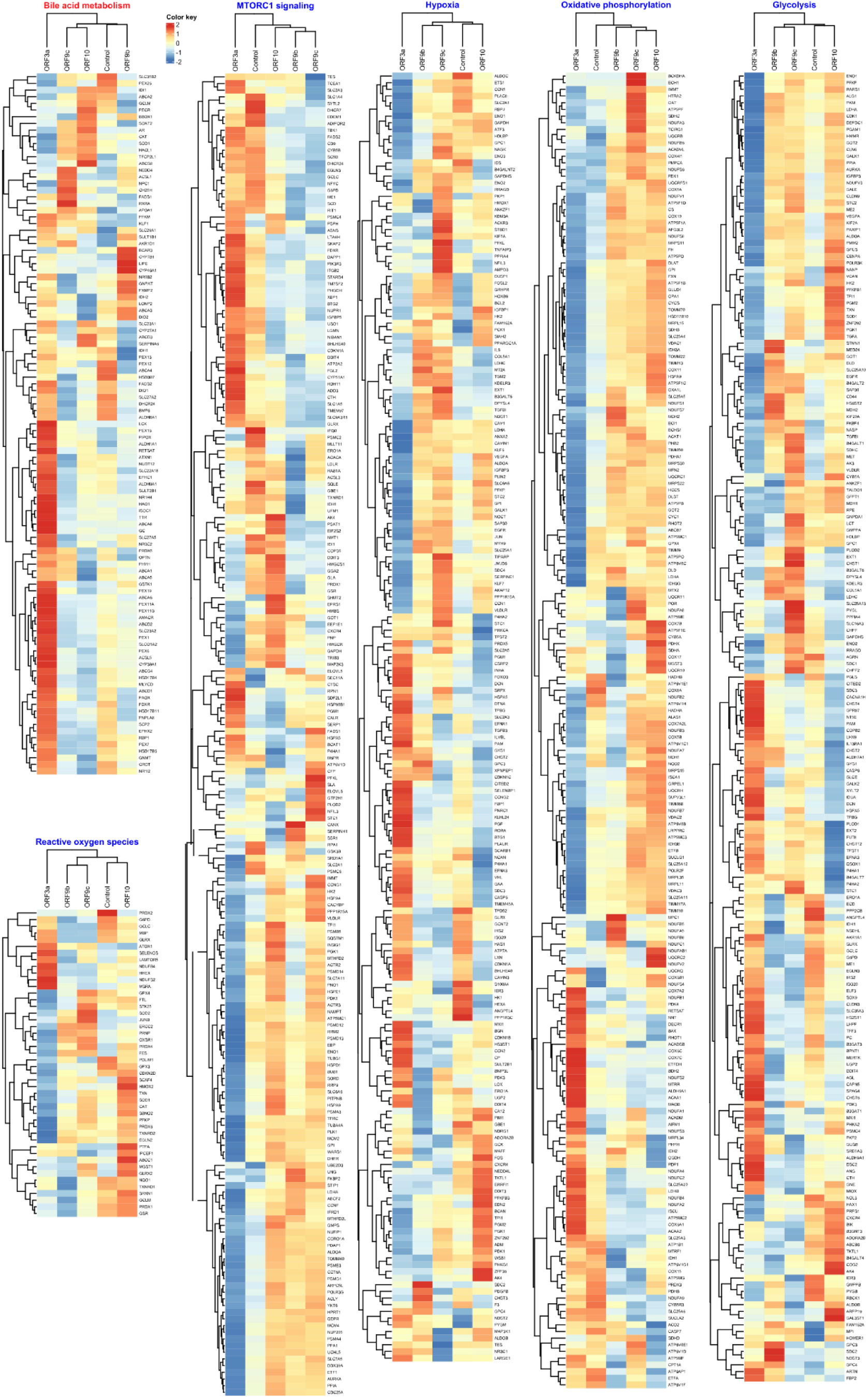
Metabolic enrichment pathways altered by ORF3a expression. The six more significant metabolic pathways up-regulated (red) and down-regulated (blue) altered by ORF3a expression are represented. Gene expression pattern of each pathway in all A549 transduced cells and A549 control cells was compared. Color key indicates Log2 FC values.

The underrepresentation of the OXPHOS pathway induced by ORF3a prompted us to conduct a more detailed examination of genes coding for mitochondrial electron transport chain (ETC) components and cristae morphology and function. GSEA with an ETC-focused gene set showed a significant underrepresentation in cells expressing ORF3a, but not the other three ORFs (Fig. 4A). This was due to the ORF3a-induced downregulation of genes for components of ETC complexes I, III, IV and V (Fig. 4A). As such, ORF3a may play a prominent role in the downregulation of mitochondrial genes and functions reported upon SARS-CoV-2 infection of lung epithelial cells^5^. By contrast, ORF9b, ORF9c and, particularly, ORF10, induced an upregulation of genes for ETC, albeit without reaching significance in terms of pathway representation by GSEA (Fig. S9). Expression analysis of genes for mitochondrial cristae morphology and function also indicated a downregulation by ORF3a of genes for critical cristae regulators, including OPA1, SAMM50, TOMM40 and components of the mitochondrial contact site and cristae organizing system (MICOS), MIC60 (IMMT) and MIC13 (MICOS13) (Fig. 4B). This is consistent with the abnormal cristae morphologies described above. Once again, ORF9b, ORF9c and ORF10 displayed transcriptional profiles for mitochondrial cristae factors components were converse to those induced by ORF3a, with upregulation of OPA1, SAMM50 or TOMM40 and downregulation of APOO (MIC26), APOOL (MIC27) or ATP5MF, particularly by ORF10 (Fig. 4B), although ATP5MF was not downregulated by ORF9c. This is also consistent with the abnormal cristae morphologies observed for ORF9b and ORF10, although likely involving mechanisms that may differ from those causing the abnormal cristae induced by ORF3a.

**Figure 4.**
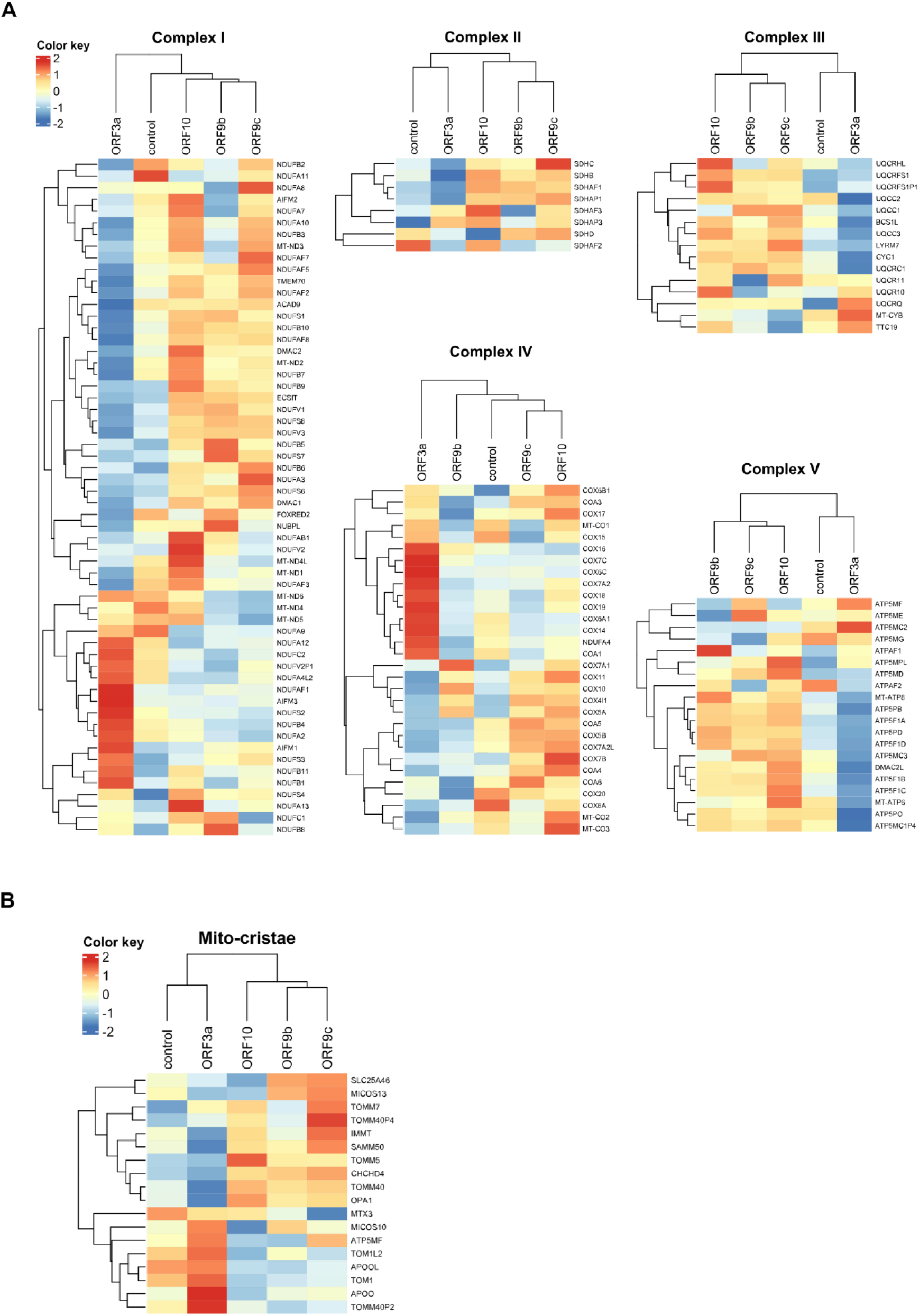
Electron Transport Chain (ETC) and mitochondrial cristae gene expression in A549 cells transduced with SARS-CoV-2 ORF3a, ORF9b, ORF9c or ORF10 accessory proteins. Heatmaps of DEGs related with mitochondrial complexes (I-V) **(A)** and mitochondria critae (Mito-cristae) **(B)** in A549 transduced cells compared with A549 control cells. Color key represents Log2 FC values.

### ORF9b overexpression causes the strongest mitochondrial function alteration

Mitochondria are a major source of ROS, and excessive production of ROS leads to oxidative damage that impairs the ability of mitochondria to make ATP and perform metabolic functions^29^. Given the above evidence of impact of the four accessory proteins on mitochondrial morphology and gene expression, ROS levels were measured with the cell-permeant H2DCFDA probe. A549 cells expressing ORF9c and ORF10 exhibited a significant increase in ROS levels compared to control cells (Fig. 5A) whereas a slight increment was detected in cells expressing ORF9b (*P*= 0.0507). In contrast, a modest reduction was observed in ORF3a-A549 cells compared to control (Fig. 5A).

**Figure 5.**
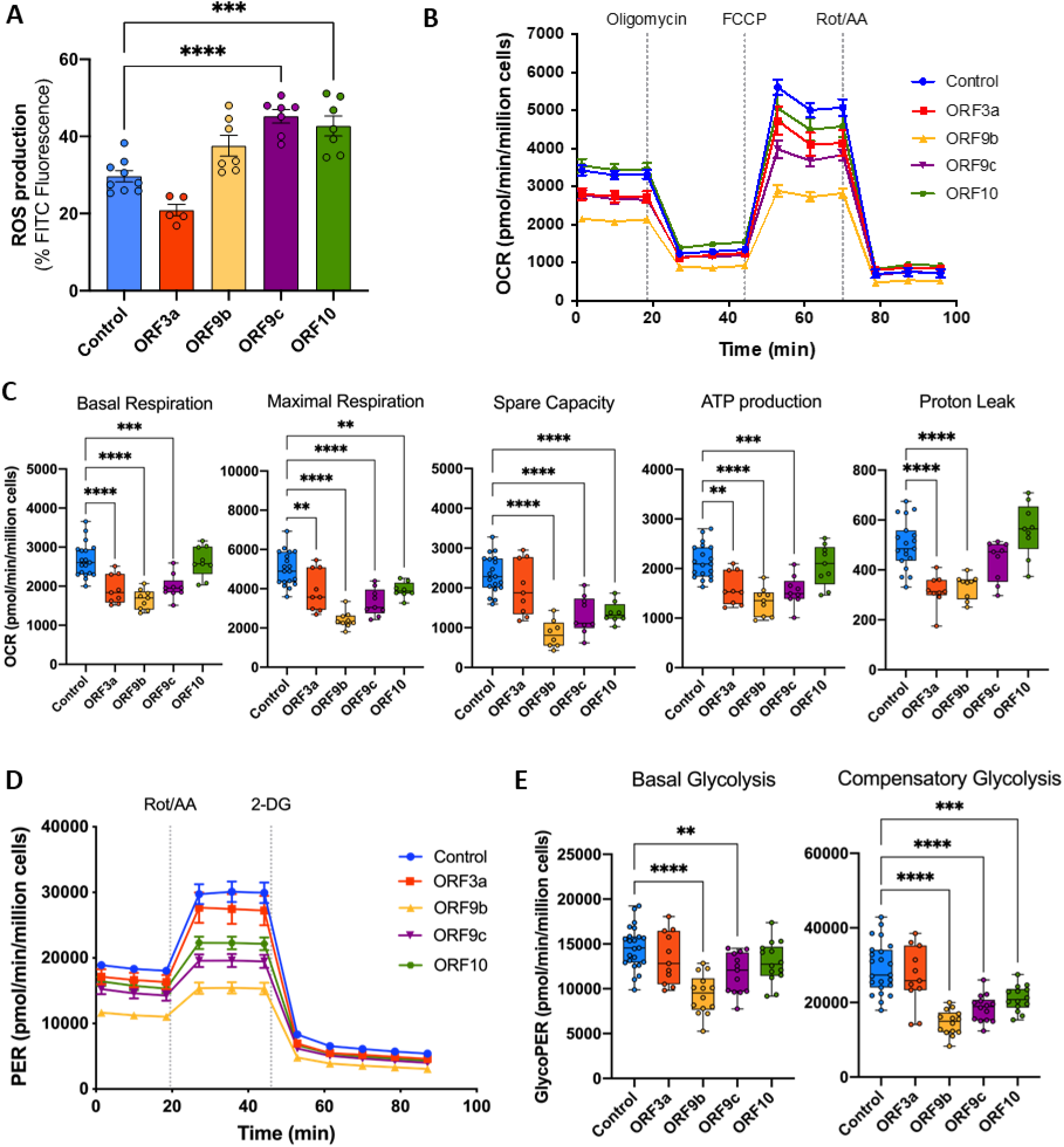
Respiratory and glycolytic profiles of A549 cells expressing SARS-CoV-2 accessory proteins. **(A)** ROS production measured by flow cytometry in A549 cells expressing SARS-CoV-2 ORF3a, ORF9b, ORF9c or ORF10 and control cells. **(B)** Seahorse XF Mito Stress test profile of oxygen consumption rate (OCR) in control and transduced cells. **(C)** Box and whisker plots showing the normalized basal respiration, maximal respiration, spare respiratory capacity, ATP production and proton leak in control and transduced cells. **(D)** Seahorse XF Glycolytic Rate test profile of proton efflux rate (PER). **(E)** Box and whisker plots showing the normalized basal and compensatory glycolytic PER. Horizontal line represents the mean and points individual replicates. Statistical significance is given as follows: *p < 0.05, **p < 0.01, ***p<0.001 and ****p<0.0001.

To determine functional consequences of SARS-CoV-2 accessory proteins on cellular bioenergetics, oxygen consumption rate (OCR), which represents the rate of oxidative phosphorylation (OXPHOS), was measured (Fig. 5B). Basal respiration, maximal respiration and ATP production were significantly impaired by ORF3a, ORF9b and ORF9c (Fig. 5C). Moreover, spare respiratory capacity was significantly hampered by ORF9b and ORF9c, but not by ORF3a. In addition, maximal respiration and spare respiratory capacity were strongly reduced by ORF9b, suggesting that this protein compromises the ability of cells to respond to high respiratory demands or to acute stress (Fig. 5C), whereas ORF10-A549 cells only showed a significant decrease in maximal and spare respiratory capacity. Proton leak, which reflects the mitochondrial oxygen consumption not coupled to ATP production, was significantly reduced by ORF3a and ORF9b, and increased by ORF10 (Fig. 5C). Additionally, glycolytic rate assays to monitor glycolysis as well as compensatory glycolysis of ORF-A549 cells were conducted. Fig. 5D shows proton efflux rate (PER) results converted from OCR and extracellular acidification rate (ECAR) data. ORF9b and ORF9c significantly reduced basal and compensatory glycolysis compared to control cells (Fig. 5E). Moreover, ORF10 also induced a decrease in compensatory glycolysis. Overall, our results indicate that all four ORFs were involved in mitochondrial dysfunction, which was particularly severe as a consequence of ORF9b expression.

### Genome-scale metabolic flux maps unveil common and private metabolic alterations induced by ORF3a, ORF9b, ORF9c and ORF10

Subsequently, metabolomics, respiration data and transcriptomics were integrated to simulate accessory protein-specific flux maps (genome-scale metabolic models, GSMM). The GIM3E^30^ and the quadratic metabolic transformation algorithms (qMTA)^31^ were applied (Supplementary table 2) for the major four metabolic superfamilies. Notably, ORF3a had a major impact on cell bioenergetics, carbohydrate metabolism and cofactors, and nucleotide metabolic pathways superfamilies (Fig. 6A).

**Figure 6.**
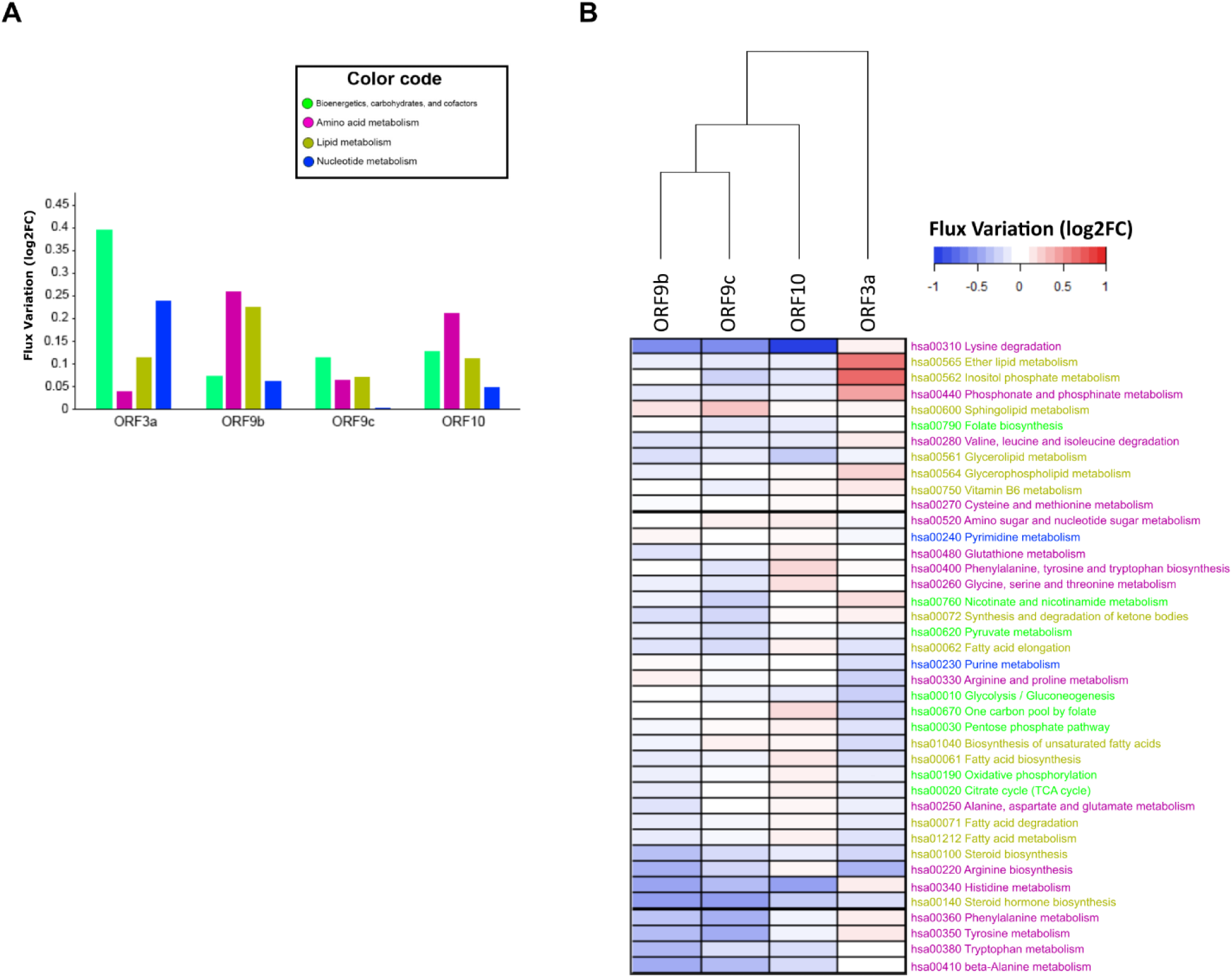
Metabolic flux changes induced by ORF3a, ORF9b, ORF9c, and ORF10 relative to control cells. **(A)** Flux variation within metabolic superfamilies. Metabolic fluxes were expressed as log2FC of ORF-A549 vs. control cells. **(B)** Heatmap representing flux modulation of ORF-A549 cells for metabolic individual KEGG pathways. Color code corresponds to metabolic superfamilies represented in **(A)**.

Detailed individual metabolic pathway analysis (Figure 6B) revealed distinctive disruptions caused by ORF3a and ORF10. Pathways uniquely increased in ORF3a-A549 cells were inositol phosphate, ether lipid, phosphonate-phosphinate, vitamin B6, glycerophospholipid metabolism and nicotinate-nicotinamide metabolism, consistent with the above-described enrichment in bile acid biosynthetic pathways and peroxisome biogenesis, where many of these processes take place ^5^, as well as essential amino acid metabolism (histidine, phenylalanine and tyrosine). Moreover, ORF3a-A549 cells displayed a distinctive decrease in fatty acid metabolism and related pathways, glycolysis, oxidative phosphorylation, metabolism of steroid hormones, arginine, proline citrate, one carbon, pentose phosphate, and purine metabolism. ORF10-A549 cells displayed a strong decrease in lysine degradation and histidine metabolism, in contrast to the slight increase observed for ORF3a-A549 cells. ORF10 also induced modest but distinctive increase in one-carbon and fatty acid metabolism-related pathways, amino acid metabolism (phenylalanine, tyrosine, tryptophan, glycine, threonine and arginine), and the citrate cycle.

ORF9b-A549 and ORF9c-A549 cells shared several metabolic communalities, including a generalized decrease in several metabolic pathways related to fatty acids, ketone bodies and amino acid metabolism, with increased sphingolipid metabolism. Interestingly, both ORFs, particularly ORF9b (Fig. 6A), induced a marked decreased in amino acid metabolism (Fig. 6B). Similarly, both ORFs had similar effects on lipid metabolism-related pathways, ORF9b causing a greater impact (Fig. 6B). Intriguingly, a common feature shared by all ORF-A549 cells was a decrease in steroid biosynthesis pathways as compared to control cells.

### Identification of putative targets to counteract the metabolic reprogramming induced by ORF3a and ORF10

Next, qMTA was used to simulate gene knock downs (KDs) that would have the potential to revert the metabolic reprogramming induced by each individual accessory protein. For each cell line, two KDs were chosen among the ten best-scoring targets (Fig. 7) predicted to be the most effective for reverting the observed metabolic phenotypes. Each outcome was linked to an existing drug, when possible. Phospholipase D2 (PLD2) and phospholipase C beta-1 (PLCB1) were the best predicted targets for reverting metabolic perturbations induced by ORF3a. PLD2 catalyzes the hydrolysis of glycerophospholipids to produce phosphatidic acid, which can be further converted into lysophosphatidic acid (LPA) and diacylglycerol (DAG). PLCB1 catalyzes the hydrolysis of 1-phosphatidylinositol 4,5-bisphosphate into DAG and inositol 1,4,5-trisphosphate (IP3). DAG and LPA are precursors for the peroxisomal and endoplasmic reticulum production of ether phospholipids^32^. Therefore, this outcome is in consonance with the above independent predictions reached by GSEA and GSMMs, indicating an upregulation of these pathways. Importantly, coronavirus envelopes are particularly enriched in ether phospholipids, along with cholesterol and sphingolipids ^33^, with a lipid composition distinct from the plasma membrane. As such, inhibiting phospholipases that catalyze the production of ether phospholipid precursors may impair the formation of viral envelopes. Interestingly, PLCB1 physically interacts with SARS-CoV-2 ORF3a^34^.

**Figure 7.**
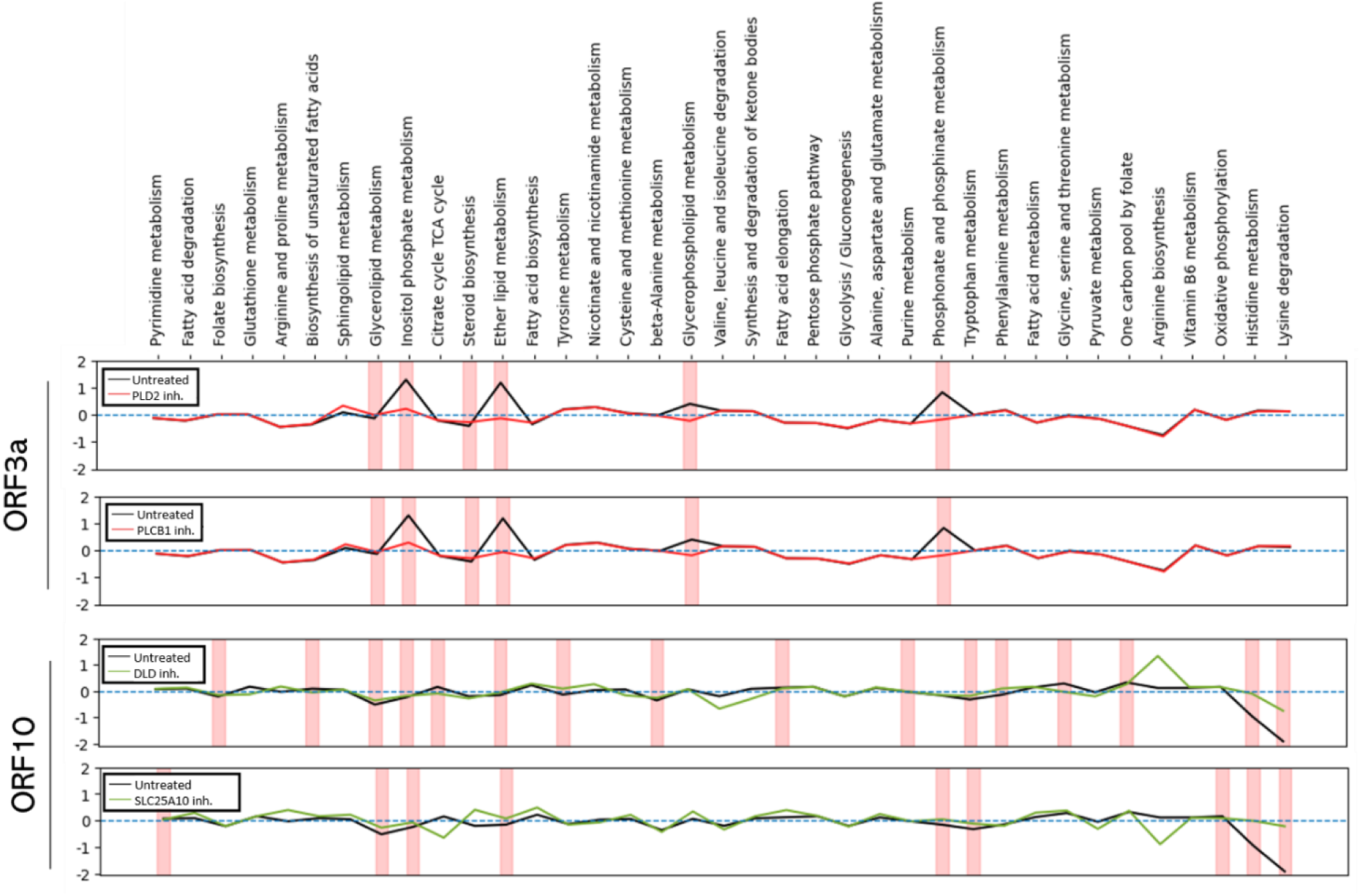
Metabolic pathway analysis representing the two-best metabolic-reversing drugs from ORF3a-A549 and ORF10-A549 cells towards the control A549 cells resting state. Each graph displays a comparison between the metabolic profile triggered by a transduction, and its reversion by *in silico* chosen drug-gene Kock downs (50% KD) selected from GSMM-mediated *in silico* analysis. Y axis represents log2FC, and the dashed-horizontal blue line at each plot represents the control state (i.e., log2FC = 0). Each line represents the metabolic change of either an ORF expression, or an ORF expression + drug, vs. control cells. PLD2: phospholipase D2; PLCB1: Phospholipase C beta-1; DLD: Dihydrolipoamide Dehydrogenase; SLC25A10: Solute Carrier Family 25 Member 10.

The best inferred targets for reverting the metabolic perturbations induced by ORF10 were dihydrolipoamide dehydrogenase (DLD), a component of the oxoglutarate dehydrogenase complex, and SLC25A10, a mitochondrion to cytosol transporter of TCA-cycle metabolites such as succinate, fumarate or malate. (Fig. 7). The inherent design of this approach, based on gene loss simulations, only allows to predict pathway perturbation reversal in association with drugs with inhibitory functions. As such, as most of the metabolic pathways perturbed by ORF9b and ORF9c showed a decreased representation (Fig. 6B), no significant drugs were predicted to revert such alterations by *in silico* simulation.

## DISCUSSION

Insights into mechanisms of SARS-CoV-2 viral life cycle and pathogenic effects are facilitated by a large number of studies of viral-host protein-protein interactions producing comprehensive interactome maps^13, 18, 35^, as well as a comprehensive GSMM of SARS-CoV-2 infected cells^2^. However, the impacts of individual accessory proteins on host cell metabolic pathways are unknown. In this study, we show that the individual expression of the SARS-CoV-2 accessory proteins, ORF3a, ORF9b, ORF9c or ORF10, in A549 lung epithelial cells results in significant mitochondrial and metabolic alterations, some of which are distinctive of a particular accessory protein. Furthermore, by combining transcriptomic analysis with functional and metabolic data in accessory protein-specific GSMMs, several alterations were identified that may point to a putative targets for investigating novel therapies. A salient outcome of our study is that, despite the distinct metabolic reprogramming caused by each of the accessory proteins studied, particularly ORF3a, all four proteins caused altered mitochondrial features, including ORF10, not known to directly associate with mitochondria.

Mitochondria are internally organized into cristae, which are fundamental structures whose shape changes under different physiological conditions^36^. The shape of the cristae is determined by the interaction of mitochondrial shape proteins^37, 38^, and perturbing such proteins disrupt cristae shape and change cristae-located OXPHOS (OXidative PHOSphorylation) system, which affects cellular growth and metabolism. Interactions between SARS-CoV-2 accessory proteins and host cell mitochondria have been proposed by multiple studies^6, 7, 17, 27^. Morphological changes in shape, size, cristae reorganization and cell location have been shown in mitochondria of SARS-CoV-2- infected cells^6, 27^. Our results show that ORF3a, ORF9c and ORF10 induce changes in the shape and density of mitochondrial cristae, suggesting that these accessory proteins may directly perturb either mitochondrial shape-determining proteins. Genes for mitochondrial proteins that determine the shape, length and function of mitochondrial cristae^37,39^, are variously downregulated by the accessory proteins studied here, particularly by ORF3a, which caused a significant downregulation of OPA1 and SAMM50. The other accessory proteins caused a downregulation of different sets of genes for proteins that also play important roles in mitochondrial and cristae morphology, suggesting that different accessory proteins disrupt mitochondrial morphology and function through different mechanisms.

All four accessory proteins induced mitochondria size reduction (mitochondria fragmentation), which was compensated by an increase in the total number of mitochondria, thus maintaining roughly equal mitochondrial masses in all cell lines. Interestingly, although ORF9b induced a significant population of smaller mitochondria (mitochondrial fragmentation or fission), the median diameter of mitochondria was larger. It is possible that a more rounded shape of the mitochondria may cause them to appear artifactually larger in longitudinal sections. Collectively, these observations are in agreement with previous studies showing mitochondrial morphological alteration and/or destabilization of normal physiological fission–fusion dynamics ensuing SARS-CoV-2 infection^6^ and suggest that such perturbations may be caused, at least partly, by these accessory proteins. Other processes, such as disruption of mitophagy as reported for ORF10^7, 17^, may also contribute to the size and shape changes in mitochondria upon viral infection, due either to excessive sequestration of healthy mitochondria or to accumulation of damaged mitochondria.

Whether a cause or a consequence of specific morphological changes in mitochondria triggered by SARS-CoV-2 and its accessory proteins studied here, they are associated with distinct metabolic and functional perturbations. As such, consistent with studies demonstrating an increased production of ROS in SARS-CoV-2 infected cells^7^, we have shown that ORF9c and ORF10 induce increased levels of ROS, which can lead to oxidative damage and abnormal energy metabolism^40^. Other disruptions to mitochondrial activity were more prominently associated with ORF9b and ORF9c, including decreased basal and maximal respiration, spare respiratory capacity, ATP production, and basal and compensatory glycolysis. Reduced basal respiration, maximal respiration, ATP production and proton leak were also associated with ORF3a expression. On the other hand, ORF10 induced reduced maximal respiration and spare respiratory capacity, as well as decreased compensatory glycolysis. Our findings of diminished bioenergetics metabolism as a caused by the four accessory proteins are consistent with the known loss of cellular energy metabolism caused by SARS-CoV-2 infection ^5, 21, 41, 42^, and are in line with the recognized interactions of ORF9b, ORF9c and ORF10 with mitochondrial factors, such as TOM70, MAVS or NIX, which may result in disrupted mitochondrial functions^12, 17^. Recently, ORF3c, another SARS-CoV-2 accessory protein, has been shown to also alter mitochondrial metabolism, inducing a shift from glucose to fatty acid oxidation and enhanced oxidative phosphorylation. ORF3c also increases ROS production and blocks the autophagic flux^43^.

The above metabolic reprogramming induced by these individual accessory proteins fails to explain the enhanced glycolysis acutely triggered by SARS-CoV-2 infection ^5^, which thus calls for a different mechanistic explanation. Applying GSEA and GSMM did not solve this issue. For example, while ORF3a-A549 cells were predicted to have a downregulation of glycolysis, experimental determination of glycolytic rates failed to show such downregulation. We speculate that compensatory mechanisms, yet to be unveiled, may regulate this process. Moreover, although our real-time functional metabolic assays demonstrated a broader range of disruptions caused by ORF9b, ORF9c and ORF10 as compared to ORF3a, GSMM predicted significant changes in energy metabolism only in association with ORF3a expression and, to a lesser degree, ORF10. In contrast to these apparently inconsistent results, other experimentally observed alterations were supported and predicted by transcriptomic and GSMM analysis, notably the correlation between altered mitochondrial cristae morphology and the downregulation of genes for critical regulators of cristae morphology and function. Morphological changes in mitochondria observed in ORF3a-A549 cells are in line with the proposed mechanism of SARS-CoV-2 replication, suggesting viral dsRNA subcellular localization and enrichment in host cell mitochondrion^6, 7^, and the alteration of the mitochondrial and endoplasmic reticulum network to viral replication organelles formed by clusters of double-membrane vesicles (DMVs)^27, 44^.

The predicted upregulation of bile acid and ether lipid biosynthesis pathways triggered by ORF3a and of one-carbon, fatty acid and amino acid metabolism induced by ORF10, led us to infer potential targets for intervention in order to revert these metabolic perturbations. A similar approach, relying on the identification of specific KDs to an existing drug, has previously revealed candidates for COVID-19 treatment ^45, 46^. In our study, qMTA simulations predicted phospholipase D2 (PLD2) and phospholipase C β-1 (PLCB1) as the best targets for reverting the metabolic phenotype induced by ORF3a in A549 cells. Of note, phosphatidic acid, the catalytic product of PLD2, has been described as essential for SARS-CoV-2 replication^47^. On the other hand, dihydrolipoamide dehydrogenase (DLD), a component of the oxoglutarate dehydrogenase complex, and SLC25A10, a mitochondrial carrier in the mitochondrial inner membrane that transports TCA-cycle metabolites such as succinate, fumarate or malate from mitochondria to cytosol, were identified as targets for reverting the ORF10-induced metabolic phenotype.

In this regard, DLD and other members of the oxoglutarate dehydrogenase complex have been reported to be decreased in SARS-CoV-2 infected Vero cells and in a cardiomyocyte model of SARS-CoV-2 infection^48^. Furthermore, in a recent multi-omics analysis in a cohort of COVID-19 patients, an essential role of transporters and metabolites of the TCA cycle has been identified in association with COVID-19 severity ^49^. Additionally, Metformin treatment, an antidiabetic-safe drug, has been recently reported to reduce the incidence of long covid. Remarkably, Metformin downregulates gene expression of SLC25A10 in A549 cells^50^.

In summary, we have found that the SARS-CoV-2 accessory proteins, ORF3a, ORF9b, ORF9c and ORF10, produce significant mitochondrial alterations and metabolic reprogramming in A549 lung epithelial cells. Although SARS-CoV-2 accessory proteins are considered non-essential for virus replication, our analyses suggest that, at least ORF3a, may have a direct effect on the viral life cycle, by coopting the host cell lipid metabolism for DMV formation, hence ultimately impacting viral replication. Finally, we have identified putative metabolic targets that could be employed to counteract the effects of SARS-CoV-2 accessory proteins ORF3a and ORF10. As such, this study illustrates GSMM as a viable tool for investigating the pathobiology of SARS-CoV-2 and as an aid to identify novel antiviral strategies.

## LIMITATIONS OF THE STUDY

This study relies on a human lung cancer cell line, with features of alveolar type 2 (AT2) lineage cells, bearing stable integration and displaying constitutive expression of a single type of viral transcript and protein per cell. We recognize that this model represents a significant departure from a situation of acute infection by SARS-CoV-2, in which the entire viral genome and proteome enter susceptible cells. Features induced or dampened by one viral protein in our reductionist models should be interpreted with caution, as downregulation of cellular transcripts or proteins by one viral protein may be compensated by the upregulation of the same transcripts or proteins elicited by a different viral protein. An advantage of such models is that it affords to gain knowledge of functions pertaining to specific viral genes that may be difficult to unravel in more complex scenarios. In recognition of this potential limitation, we have placed particular emphasis on the analysis of features upregulated, rather than downregulated by viral accessory proteins, as they should contribute to phenotypes and perturbations induced also by complex virions, independent of downregulation by distinct viral factors. A further limitation, beyond complex vs. reductionist approaches, stems from the long-term expression of viral proteins in our cell models, as opposed to acute viral infections, which involve short-term virus-cell interactions. The analysis of prolonged interactions reflects indirect cellular adaptations to long-term perturbations, in addition to more direct effects by the viral protein under study. However, it could be interested for further investigations in long COVID-19 treatments. Finally, we identify metabolic changes observed upon ORF transduction, but such alterations do not necessarily need to be beneficial for infection and viral replication and might instead be cellular responses which aim to protect against ORFs induced cellular damage. For instance, SOD activity might protect against SARS-CoV-2 induced oxidative stress^51^.

## MATERIAL & METHODS

### Cell culture, lentivirus production and transduction

A549 pulmonary epithelial cells (ATCC CRM-CCL-185; RRID: CVCL_0023) were cultured in Dulbecco’s Modified Eagle Medium (DMEM) (Gibco, #41966029) supplemented with 10% (v/v) heat-inactivated fetal bovine serum (FBS) (Gibco, #1027016), 1% Penicillin-Streptomycin (100U/ml) (Gibco, #15070063) and Amphotericin B (Gibco, #15290026). ORF3a, ORF9b, ORF9c or ORF10 accessory proteins coding sequences (codon-optimized for mammalian expression) were cloned into pLVX-EF1α-IRES-Puro Cloning and Expression Lentivector (Clontech, Takara, #631253) to generate pseudotyped lentiviral particles encoding each accessory protein of SARS-CoV-2 (Wuhan-Hu-1 isolate) at the CNIC (Centro Nacional de Investigaciones Cardiovasculares) Viral Vector Unit (ViVU) as described previously^19, 52^. Accessory proteins were C-terminally 2xStrep-tagged to check viral protein expression. A549 cells were transduced by incubating them with lentivirus at a MOI of 10 for 24 h followed by 2 µg/ml puromycin treatment to start the selection of successfully transduced cells. All cells were cultured at 37°C in a 5% CO2, 90% humidity atmosphere.

### Strep-tag Immunofluorescence, RNA isolation and sequencing

These methods were performed as previously described^19, 52^.

### Mitochondria Analysis

#### Transmission Electron Microscopy (Ultrastructural Analysis of Cells)

Cell monolayers were washed with PBS 1x and fixed *in situ* for 1 hour at room temperature (RT) with 3 % glutaraldehyde (EM Grade, Ted Pella INC) in PBS 1x. Fixed cells were washed three times with PBS 1x. Cell post-fixation was as follows: 1 hour at 4°C with 1% osmium tetroxide (Electron Microscopy Sciences) and 0,8% potassium ferricyanide in PBS, washed with PBS three times, and after dehydration in a gradient of 30% to 100% in ETOH, cells were embedded in a gradient of EtOH/ LX 112 epoxy resin to 100% epoxy resin (Ladd Research). The samples were polymerized at 60°C for two days. Ultrathin 70 nm-thick sections were obtained with a Leica EM UC6 ultramicrotome (Leica Microsystems GmbH), transferred to collodion/carbon-coated EM grids and stained with Uranyl Acetate 5% for 20 min and Reynold´s Lead Citrate for 5 min. 70 nm sections were visualized on a FEI Tecnai 12 electron microscope equipped with a LaB6 filament and operated at 100 kV. Images were recorded with a FEI Ceta digital camera at various magnifications. Median of mitochondria longest diameter was measured with ImageJ program, and median of ridges was counted in high resolution ME images. In both cases, at least 50 mitochondria were analyzed.

#### TOM20 immunofluorescence

12,000 cells per well were seeded on a chamber slide with a removable 12 well chamber (ibidi, #81201). Cells were fixed with 4% PFA in PBS for 5 min, washed three times in PBS, and then permeabilized for 10 min with 0.1% Triton X-100 in PBS. TOM20 (abcam, ab186735) incubation was carried out for 1h in PBS containing 3% BSA and 0.1% Triton X-100 at 1:100 dilution. Chamber slides were washed three times with PBS before Alexa 488 secondary anti-mouse antibody (Jackson, #155-546-062) incubation (1:1000 dilution). Texas Red-X phalloidin (Invitrogen, #T7471) was used as a cytoplasmic marker at 1:400, and DAPI (4’6-diamidino-2- phenylindole) (Thermo Fisher Scientific, #62248) was used as a nuclear marker at a final concentration of 1µg/ml. After removing the chamber, the slides were mounted with FluorSave^TM^ Reagent (Merck, #345789) and coverslips of 1.5H thickness were used.

#### Optical microscopy

Images were acquired with a Thunder Imager Microscope (Leica Microsystems). To ensure reliable cell comparisons, only single cells were acquired; this avoids differences due to cell-cell adhesion and reduces segmentation errors. To identify single cells, we used a custom-made program named AIMS. This program uses, first, a fast-low-resolution scan (5X) to detect cells (Phalloidin - Actin) and nuclei (DAPI-DNA), and, then, an image analysis macro runs discarding objects bigger or smaller than a single cell. After this, all the coordinates for single cells localized in AIMS are loaded and acquired automatically in Thunder imager (40X HC PL APO 0,95 NA dry objective).

#### Image analysis

To identify mitochondrial size (mitochondria fragmentation), number per cell, and area covered by mitochondria (mitochondrial mass), at least 200 cells were analyzed. Images were processed by using Cell profiler^53^. Statistical analysis was performed with SPSS v19.0 (IBM).

#### Tracking analysis

To estimate the motility of mitochondria, 50.000 or 75.000 cells per well were seeded on a chamber slide (ibidi, #80826) and stained with MitoTracker Red. The emitted fluorescence was imaged using Thunder imager (40X HC PL APO 0,95 NA dry objective), time lapse data were recorded at 33 frame per second and field of views were randomly selected from the imaging software (Leica) across the sample. The tracking analysis was performed with TrackMate 7^54^. Due to their non-circular shape, mitochondria were identified using a mask detector. The masks were generated for each frame using a custom script. Here, we subtracted the background light in each frame via top hat filtering and evened out the fluorescence signal with a high-pass filter to avoid the misdetection of dimmer organelles at the cell borders. Finally, to reject possible lingering background signal, we generated the masks using an adaptive threshold calculated considering the local background levels. Afterwards, mitochondria motion and speed were estimated using a sparse LAP tracker^55^. To obtain more robust estimate of the average speed of each detected track, we only considered tracks with at least 20 time points.

#### Flow cytometry

Cells were seeded at 3 x 10^5^ cells/well in 6-well plates 24 h prior to staining. For quantitative analysis of membrane potential, cells were stained in DMEM 1% penicillin/streptomycin with 5 nM MitoView 533 (Biotium, #70055). Plates were incubated at 37°C for 30 min protected from light. Cells were then harvested, washed with PBS 1X and resuspended in PBS. For these experiments, 10.000 events were analysed using a CytoFLEX flow cytometer (Beckman Coulter) and FlowJo v10 software (BD Biosciences).

### Differential gene expression

The R/Bioconductor package DESeq2 v.1.40.2^56^ was used to evaluate the differential expression among the experimental groups (control vs ORF). Genes for which the sum of raw counts across all of the samples was < 10 were discarded. Genes with an adjusted p-value ≤ 0.05 and an absolute log_2_ fold change ≥ 1 were considered differentially expressed. The R package pheatmap v.1.0.12 (pheatmap: Pretty Heatmaps_. R package version 1.0.12) was used to generate the heatmaps of gene expression values obtained with DESeq2, normalized by variance stabilizing transformation (vst) and by rows.

### Gene Set Enrichment Analysis ^26^

GSEAPreranked was used to assess gene enrichment compared to gene sets from the MSigDB collection (Hallmark gene sets^57^). A separate GSEA was performed with gene sets for electron transport chain components^5^ and mitochondria cristae morphology and function^58^. GSEA was performed with 1,000 random permutations to yield FDR q-values and normalized enrichment scores (NES). Genes were ranked following the Wald statistic obtained with DESeq2.

### Reactive oxygen species (ROS) production analysis

ROS production was evaluated using the ROS Detection Assay Kit (Canvax Biotec S.L, Cordoba, Spain) which contains the cell-permeant reagent dichlorodihydrofluorescein diacetate (H_2_DCFDA), an indicator of reactive oxygen intermediates, that becomes fluorescent when oxidized, following the manufacturer’s instructions. Cells were seeded on a 24 well plates at 7,5×10^4^ cells per well 2-day prior assay. Cells were incubated with H_2_DCFDA (25 mM) for 45 minutes at 37°C. Positive control cells were treated with 100 µM H_2_O_2_. ROS were measured by flow cytometry (BD Accuri C6 Plus Flow Cytometer).

### Seahorse extracellular flux analysis

Agilent Seahorse XF Cell Mito Stress Test was applied, and oxygen consumption rate (OCR) and extracellular acidification rate (ECAR) were determined using Agilent Seahorse XF24 Analyzer (Seahorse Bioscience, Agilent) previously calibrated using Seahorse XF Calibrant solution in a CO_2_-free incubator overnight. 24 h before the assay, cells were seeded at 37,500 cells per well in a Seahorse 24-well XF Cell Culture microplate in DMEM culture medium and were allowed to adhere for 24 h in 5% CO_2_ atmosphere at a 37 °C. On the day of assay, media was changed to XF DMEM medium supplemented with 10 mM glucose, 1 mM sodium pyruvate and 2 mM glutamine, pH 7.4, and then maintained in XF assay media at 37 °C in a CO_2_-free incubator 1 h. Mitochondrial function of the cells was analyzed by sequential injections of the modulators oligomycin (1 μM), carbonyl cyanide-*p*- trifluoromethoxyphenylhydrazone (FCCP, 1 μM) and a mixture of antimycin A and rotenone (Rot/AA, 0.5 μM).

For glycolytic rate analysis, the medium was changed to Seahorse XF DMEM medium (10 mM glucose, 1 mM sodium pyruvate, 2 mM glutamine, pH 7.4). Then, ECAR and OCR were recorded using the Agilent Seahorse XF24 Analyzer following injections with 0.5 µM Rot/AA and 50 mM 2-deoxyglucose (2-DG) (XF Glycolytic Rate Test Kit), respectively. Data were collected using Agilent Seahorse Wave 2.6.1 Desktop software and normalized to protein concentration determined at the end of the assay. Data were exported to GraphPad Prism version 9 for analysis.

### Measurement of consumption and production rates of metabolites

For measurement of uptake and production rates of metabolites, 2 x10^5^ cells per well were seeded in triplicate in 6-well plates with standard incubation medium. After 24h, cells were counted and supernatants were collected and frozen until further analysis. Consumption and production rates of metabolites were determined by measuring metabolite concentration in incubation media at the beginning and at the end of 24h-incubation time and correcting the absolute consumption/production by time and cell number assuming lineal cell growth.

Glucose, lactate, glutamate and glutamine concentrations in medium aliquots were determined spectrophotometrically using NAD(P)H-coupled enzymatic reactions in an autoanalyzer Cobas Mira Plus (Horiba ABX, Kyoto, Japan)^59^. Concentration of 180 metabolites from 6 different families (amino acids, biogenic amines, acilcarnitines, glycerophospholipids, esphingolipids and hexoses) were determined using the Absolute IDQ p180 kit from Biocrates Life Sciences (Innsbruck, Austria) and an AB Sciex 6500 QTRAP MS/MS mass spectrometer coupled to a UHPLC Agilent 1290 Infinity II System. 10 μl of media was plated in each well, and the kit was processed following manufacturer’s instructions. Analyst and the MetIDQ™ software packages were used to analyse the obtained data and calculate metabolite concentrations^59^.

### Construction of condition-specific GSMMs

Condition-specific Genome-Scale Metabolic Models (GSMMs) of control and ORF-A549 cells were reconstructed using the human GSMM Recon3D^60^ as a template by integrating transcriptomics (RNAseq), uptake and production rates of metabolites (measured as previously indicated) and Seahorse extracellular fluxes, using the COBRApy toolbox^61^. As part of this reconstruction, enzymes with low expression were removed from the network. More in detail, enzymes with average gene expression (FPKM) value under 1 in all conditions were removed, always ensuring their removal still allowed to produce 50% of optimal biomass and to sustain the measured rates of metabolite uptake and secretion. In addition, enzymes with FPKM values under 1 in any of the ORF-A549 cell lines, were also removed from the condition-specific model if the difference against the control was statistically significant (FDR < 0.05).

### Quadratic metabolic transformation algorithm

The metabolic transitions of ORF-A549 when were simulated with the quadratic metabolic transformation algorithm (qMTA)^31^. qMTA was run using as an input the condition-specific models, the reference (i.e., control) flux distribution for A549 cells and the gene expression and measured rates fold-change between control cells and ORF-A549 cells (measured rates shown in Supplementary Table 2). The reference flux distribution was defined as the average flux values when sampling the solution space within 99% of GIME3’s optimal solution^30, 31^ in the A549 control model (i.e., non-transduced A549 cells).

Briefly, qMTA simulates the metabolic transitions of A549 when transduced with ORFs by maximizing the consistency between the variation in gene expression and measured rates and the resulting flux distribution relative to the reference flux distribution. As part of this process p-values for differential gene expression and standard deviation for measured flux rates are used to give more weight measured variations with a stronger statistical significance. The result is a flux distribution consistent with the transcriptomics and measured rates for each transfection, i.e., ORF3a, ORF9b, ORF9c, and ORF10 (ORF-A549 cells).

To facilitate summarizing the results of the metabolic transformation, individual fluxes are assigned to a KEGG pathway. This assignment is achieved by matching the genes catalyzing each reaction to the gene-pathway annotation of KEGG. Then, the individual reaction fluxes of each pathway are added (in absolute value) to compute the total flux value for each pathway. To further summarize the results, KEGG pathways were also grouped to four metabolic pathway superfamilies (Bioenergetics and carbohydrate, amino acids, lipids and nucleotide metabolism) and total flux value for each superfamily was calculated using the same procedure.

### Identification of targets disrupting metabolic transformations

qMTA is also used to identify putative gene knock downs and metabolic inhibitors that can disrupt the metabolic transformation underlying ORF transduction. This is achieved for each condition by iteratively repeating the qMTA analysis with gene knock downs (simulated by reducing the maximum flux to each mapped reaction to 50% of the control condition). This tests the capacity of a gene knock down to prevent or disrupt the metabolic transformation. To increase robustness^62^, this analysis is complemented by using the minimization of metabolic adjustment (MOMA) algorithm^63^ to simulate the capacity of individual gene knock downs to switch the ORF metabolic flux distribution to a state closer to the A549 control cells. Briefly, MOMA simulates the effect of a gene knock down by reducing the maximum flux to each reaction mapped to gene to 50% and minimizing the variation in the remaining reactions. For each condition, putative genes knock downs are ranked based on their capacity to both disrupt the metabolic transformation from control to ORF-A549 cells and their capacity to revert the ORF-transduced state to the control.

## DATA AVAILABILITY

The datasets generated during the current study are available in the NCBI BioProject database (https://www.ncbi.nlm.nih.gov/bioproject/) under accession number PRJNA975806 for ORF3a-A549 cells, PRJNA946640 for ORF9b and ORF9c transduced cells, PRJNA976185 for ORF10-A549 cells, and PRJNA841835 for A549 control cells.

## AUTHOR CONTRIBUTIONS

Conceptualization: MM

Methodology: all authors

Investigation: all authors

Funding acquisition: MM, JJG, MC

Writing – original draft: MM, BD, JG, MC and TT

Writing – review & editing: MM, BD

## REFERENCES

1. World Health Organization. WHO Coronavirus (COVID-19) Dashboard.) (2023, March 21).

2. Cheng K, et al. Genome-scale metabolic modeling reveals SARS-CoV-2-induced metabolic changes and antiviral targets. Molecular Systems Biology 17, (2021).

3. Pei R, et al. Host metabolism dysregulation and cell tropism identification in human airway and alveolar organoids upon SARS-CoV-2 infection. Protein Cell 12, 717–733 (2021).

4. Gardinassi LG, Souza COS, Sales-Campos H, Fonseca SG. Immune and Metabolic Signatures of COVID-19 Revealed by Transcriptomics Data Reuse. Frontiers in Immunology 11, (2020).

5. Guarnieri JW, et al. Core mitochondrial genes are down-regulated during SARS-CoV-2 infection of rodent and human hosts. Sci Transl Med 15, eabq1533 (2023).

6. Bhowal C, Ghosh S, Ghatak D, De R. Pathophysiological involvement of host mitochondria in SARS-CoV-2 infection that causes COVID-19: a comprehensive evidential insight. Molecular and Cellular Biochemistry, (2022).

7. Shang C, et al. SARS-CoV-2 Causes Mitochondrial Dysfunction and Mitophagy Impairment. Front Microbiol 12, 780768 (2021).

8. West AP, Shadel GS, Ghosh S. Mitochondria in innate immune responses. Nature Reviews Immunology 11, 389–402 (2011).

9. Burtscher J, Cappellano G, Omori A, Koshiba T, Millet GP. Mitochondria: In the Cross Fire of SARS-CoV-2 and Immunity. iScience 23, 101631 (2020).

10. Redondo N, Zaldívar-López S, Garrido JJ, Montoya M. SARS-CoV-2 Accessory Proteins in Viral Pathogenesis: Knowns and Unknowns. Front Immunol 12, 708264 (2021).

11. Hurtado-Tamayo J, Requena-Platek R, Enjuanes L, Bello-Perez M, Sola I. Contribution to pathogenesis of accessory proteins of deadly human coronaviruses. Front Cell Infect Microbiol 13, 1166839 (2023).

12. Gao X, Zhu K, Qin B, Olieric V, Wang M, Cui S. Crystal structure of SARS-CoV-2 Orf9b in complex with human TOM70 suggests unusual virus-host interactions. Nature Communications 12, (2021).

13. Zhou Y, et al. A comprehensive SARS-CoV-2–human protein–protein interactome reveals COVID-19 pathobiology and potential host therapeutic targets. Nature Biotechnology 41, 128–139 (2023).

14. Ren Y, et al. The ORF3a protein of SARS-CoV-2 induces apoptosis in cells. Cellular & Molecular Immunology 17, 881–883 (2020).

15. Wang R, et al. ORF3a Protein of Severe Acute Respiratory Syndrome Coronavirus 2 Inhibits Interferon-Activated Janus Kinase/Signal Transducer and Activator of Transcription Signaling via Elevating Suppressor of Cytokine Signaling 1. Front Microbiol 12, 752597 (2021).

16. Miller AN, et al. The SARS-CoV-2 accessory protein Orf3a is not an ion channel, but does interact with trafficking proteins. eLife 12, (2023).

17. Li X, et al. SARS-CoV-2 ORF10 suppresses the antiviral innate immune response by degrading MAVS through mitophagy. Cellular & Molecular Immunology 19, 67–78 (2022).

18. Gordon DE, et al. A SARS-CoV-2 protein interaction map reveals targets for drug repurposing. Nature 583, 459–468 (2020).

19. Lopez-Ayllon BD, et al. SARS-CoV-2 accessory proteins involvement in inflammatory and profibrotic processes through IL11 signaling. Front Immunol 14, 1220306 (2023).

20. Miserey-Lenkei S, et al. A comprehensive library of fluorescent constructs of SARS-CoV-2 proteins and their initial characterisation in different cell types. Biol Cell 113, 311–328 (2021).

21. Ramachandran K, et al. SARS-CoV-2 infection enhances mitochondrial PTP complex activity to perturb cardiac energetics. iScience 25, 103722 (2022).

22. Jeyaraju DV, Cisbani G, Pellegrini L. Calcium regulation of mitochondria motility and morphology. Biochim Biophys Acta 1787, 1363–1373 (2009).

23. Miller KE, Sheetz MP. Axonal mitochondrial transport and potential are correlated. J Cell Sci 117, 2791–2804 (2004).

24. Dominguez Andres A, et al. SARS-CoV-2 ORF9c Is a Membrane-Associated Protein that Suppresses Antiviral Responses in Cells. bioRxiv, (2020).

25. Han L, et al. SARS-CoV-2 ORF10 antagonizes STING-dependent interferon activation and autophagy. J Med Virol 94, 5174–5188 (2022).

26. Subramanian A, et al. Gene set enrichment analysis: a knowledge-based approach for interpreting genome-wide expression profiles. Proc Natl Acad Sci U S A 102, 15545–15550 (2005).

27. Cortese M, et al. Integrative Imaging Reveals SARS-CoV-2-Induced Reshaping of Subcellular Morphologies. Cell Host Microbe 28, 853–866 e855 (2020).

28. Ferreira V, Ferreira AR, Ribeiro D. Peroxisomes and Viruses: Overview on Current Knowledge and Experimental Approaches. Methods Mol Biol 2643, 271–294 (2023).

29. Venditti P, Di Meo S. The Role of Reactive Oxygen Species in the Life Cycle of the Mitochondrion. Int J Mol Sci 21, (2020).

30. Schmidt BJ, Ebrahim A, Metz TO, Adkins JN, Palsson B, Hyduke DR. GIM3E: condition-specific models of cellular metabolism developed from metabolomics and expression data. Bioinformatics 29, 2900–2908 (2013).

31. Foguet C, et al. Genetically personalised organ-specific metabolic models in health and disease. Nat Commun 13, 7356 (2022).

32. Dean JM, Lodhi IJ. Structural and functional roles of ether lipids. Protein Cell 9, 196–206 (2018).

33. Saud Z, et al. The SARS-CoV2 envelope differs from host cells, exposes procoagulant lipids, and is disrupted in vivo by oral rinses. J Lipid Res 63, 100208 (2022).

34. http://www.sars-cov-2-interactome.org.

35. Alqutami F, Senok A, Hachim M. COVID-19 Transcriptomic Atlas: A Comprehensive Analysis of COVID-19 Related Transcriptomics Datasets. Frontiers in Genetics 12, (2021).

36. Cogliati S, et al. Mitochondrial cristae shape determines respiratory chain supercomplexes assembly and respiratory efficiency. Cell 155, 160–171 (2013).

37. Frezza C, et al. OPA1 controls apoptotic cristae remodeling independently from mitochondrial fusion. Cell 126, 177–189 (2006).

38. Pfanner N, et al. Uniform nomenclature for the mitochondrial contact site and cristae organizing system. J Cell Biol 204, 1083–1086 (2014).

39. Ott C, et al. Sam50 functions in mitochondrial intermembrane space bridging and biogenesis of respiratory complexes. Mol Cell Biol 32, 1173–1188 (2012).

40. Zorov DB, Juhaszova M, Sollott SJ. Mitochondrial reactive oxygen species (ROS) and ROS-induced ROS release. Physiol Rev 94, 909–950 (2014).

41. Saleh J, Peyssonnaux C, Singh KK, Edeas M. Mitochondria and microbiota dysfunction in COVID-19 pathogenesis. Mitochondrion 54, 1–7 (2020).

42. Singh KK, Chaubey G, Chen JY, Suravajhala P. Decoding SARS-CoV-2 hijacking of host mitochondria in COVID-19 pathogenesis. Am J Physiol Cell Physiol 319, C258–C267 (2020).

43. Mozzi A, et al. SARS-CoV-2 ORF3c impairs mitochondrial respiratory metabolism, oxidative stress, and autophagic flux. iScience 26, 107118 (2023).

44. Wolff G, et al. A molecular pore spans the double membrane of the coronavirus replication organelle. Science 369, 1395–1398 (2020).

45. Su J, et al. SARS-CoV-2 ORF3a inhibits cGAS-STING-mediated autophagy flux and antiviral function. J Med Virol 95, e28175 (2023).

46. Krishnakumar R, et al. FOXD3 Regulates Pluripotent Stem Cell Potential by Simultaneously Initiating and Repressing Enhancer Activity. Cell Stem Cell 18, 104–117 (2016).

47. Tabata K, et al. Convergent use of phosphatidic acid for hepatitis C virus and SARS-CoV-2 replication organelle formation. Nat Commun 12, 7276 (2021).

48. Mullen PJ, et al. SARS-CoV-2 infection rewires host cell metabolism and is potentially susceptible to mTORC1 inhibition. Nat Commun 12, 1876 (2021).

49. Ambikan AT, et al. Multi-omics personalized network analyses highlight progressive disruption of central metabolism associated with COVID-19 severity. Cell Syst 13, 665–681 e664 (2022).

50. Zhao Q, Zhou X, Curbo S, Karlsson A. Metformin downregulates the mitochondrial carrier SLC25A10 in a glucose dependent manner. Biochem Pharmacol 156, 444–450 (2018).

51. Mo LH, et al. Superoxide Dismutase Prevents SARS-CoV-2-Induced Plasma Cell Apoptosis and Stabilizes Specific Antibody Induction. Oxid Med Cell Longev 2022, 5397733 (2022).

52. Garcia-Garcia T, et al. Impairment of antiviral immune response and disruption of cellular functions by SARS-CoV-2 ORF7a and ORF7b. iScience 25, 105444 (2022).

53. Stirling DR, Swain-Bowden MJ, Lucas AM, Carpenter AE, Cimini BA, Goodman A. CellProfiler 4: improvements in speed, utility and usability. BMC Bioinformatics 22, 433 (2021).

54. Ershov D, et al. TrackMate 7: integrating state-of-the-art segmentation algorithms into tracking pipelines. Nat Methods 19, 829–832 (2022).

55. Jaqaman K, et al. Robust single-particle tracking in live-cell time-lapse sequences. Nat Methods 5, 695–702 (2008).

56. Love MI, Huber W, Anders S. Moderated estimation of fold change and dispersion for RNA-seq data with DESeq2. Genome Biol 15, 550 (2014).

57. Liberzon A, Birger C, Thorvaldsdottir H, Ghandi M, Mesirov JP, Tamayo P. The Molecular Signatures Database (MSigDB) hallmark gene set collection. Cell Syst 1, 417–425 (2015).

58. Giacomello M, Pyakurel A, Glytsou C, Scorrano L. The cell biology of mitochondrial membrane dynamics. Nat Rev Mol Cell Biol 21, 204–224 (2020).

59. de Atauri P, et al. Carbon metabolism and the sign of control coefficients in metabolic adaptations underlying K-ras transformation. Biochim Biophys Acta 1807, 746–754 (2011).

60. Brunk E, et al. Recon3D enables a three-dimensional view of gene variation in human metabolism. Nat Biotechnol 36, 272–281 (2018).

61. Ebrahim A, Lerman JA, Palsson BO, Hyduke DR. COBRApy: COnstraints-Based Reconstruction and Analysis for Python. BMC Syst Biol 7, 74 (2013).

62. Valcárcel LV, Torrano V, Tobalina L, Carracedo A, Planes FJ. rMTA: robust metabolic transformation analysis. Bioinformatics 35, 4350–4355 (2019).

63. Segrè D, Vitkup D, Church GM. Analysis of optimality in natural and perturbed metabolic networks. Proc Natl Acad Sci U S A 99, 15112–15117 (2002).

