## Supplementary Figures for "Metabolic and mitochondria alterations induced by SARS-CoV-2 accessory proteins ORF3a, ORF9b, ORF9c and ORF10"

#### Supplementary Figure 1

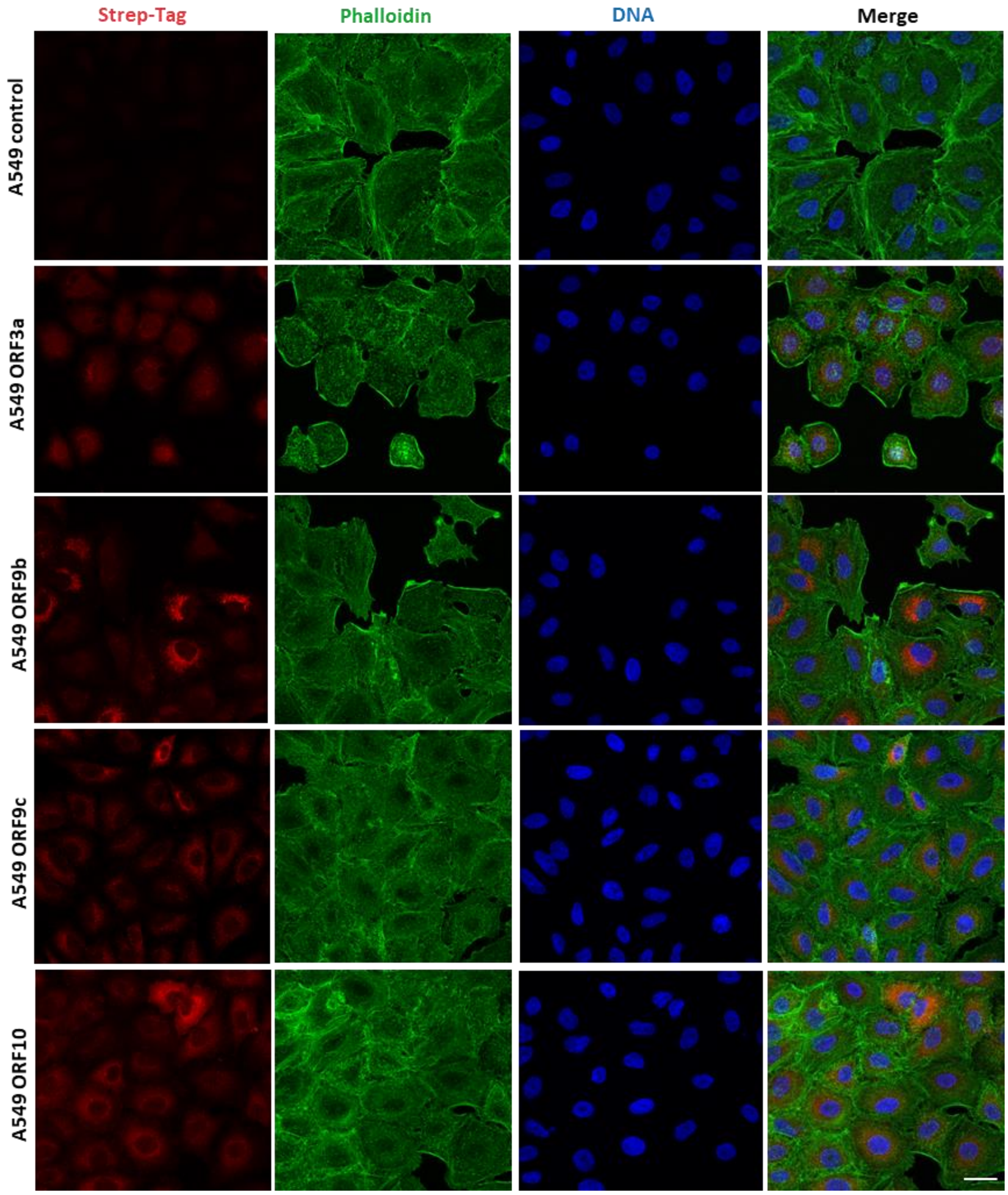


**Figure S1.** Representative confocal images of A549 cells expressing SARS-CoV-2 ORF3a, ORF9b, ORF9c or ORF10 (objective 63x, scale bar 25 µm).

#### Supplementary Figure 2

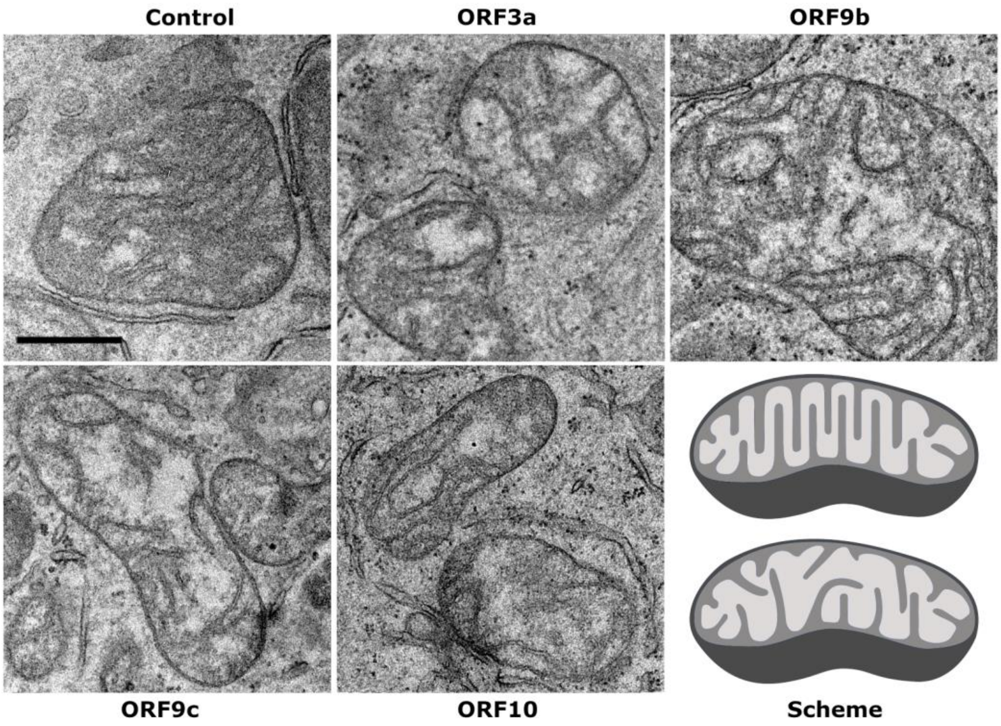


**Figure S2**. Mitochondrial morphology in ORF-A549 cells. Analysis by transmission electron microscopy of mitochondrial appearance in ORF-A549 cells ultrathin sections. Representative images of mitochondria in each condition. Scale bar indicates 500 nm. Last panel shows a schematic representation of mitochondria with organized and parallel mitochondrial ridges (top), and mitochondria with disorganized, non-parallel mitochondrial ridges and hollowed out areas (bottom).

#### Supplementary Figure 3

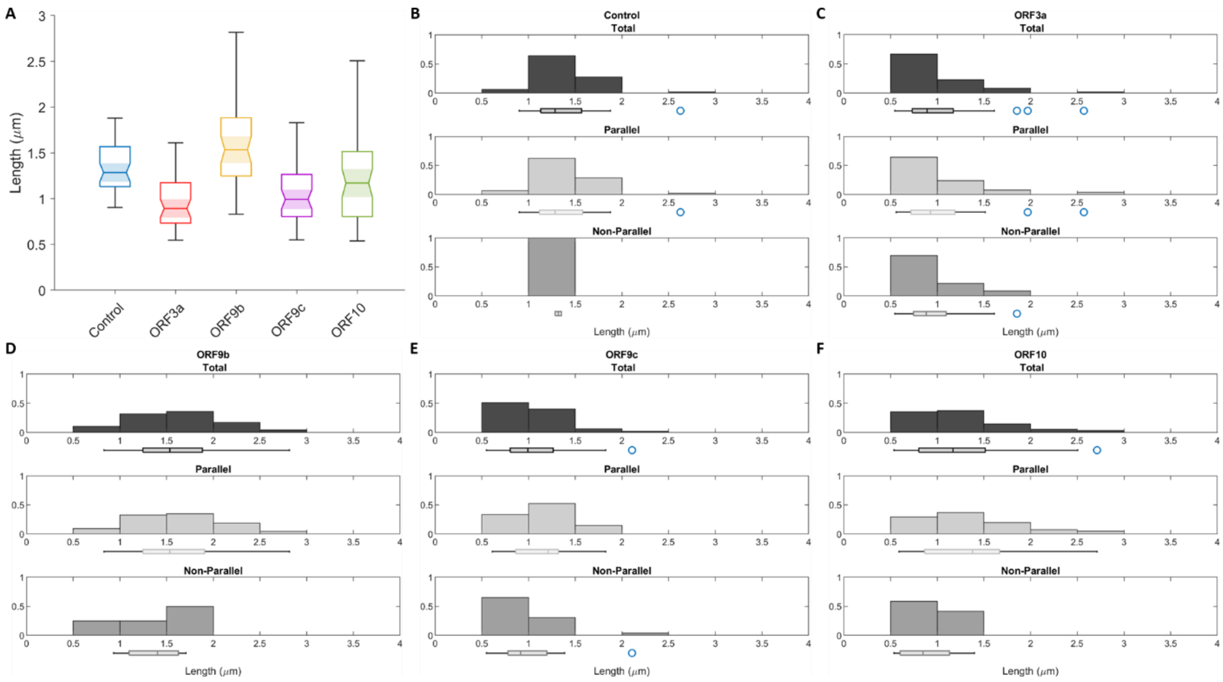


**Figure S3.** Distribution of the length of mitochondria in ORF-A549 cells imaged by transmission electron microscopy. **(A)** Box chart summarizing the distribution of the mitochondrial length measured in each cell line. Each box is divided into four sections representing a different quartile while the central lines are the median values and the areas marked in solid colors are the intervals of significance. The full distributions have been depicted in the histograms in **(B-F)**. For each cell line, the corresponding distribution of lengths has been plotted considering the total amount of measurements (upper histograms), as well as separating the mitochondria showing only parallel ridges from the mitochondria with non-parallel ones (middle and lower histograms).

### Supplementary Figure 4

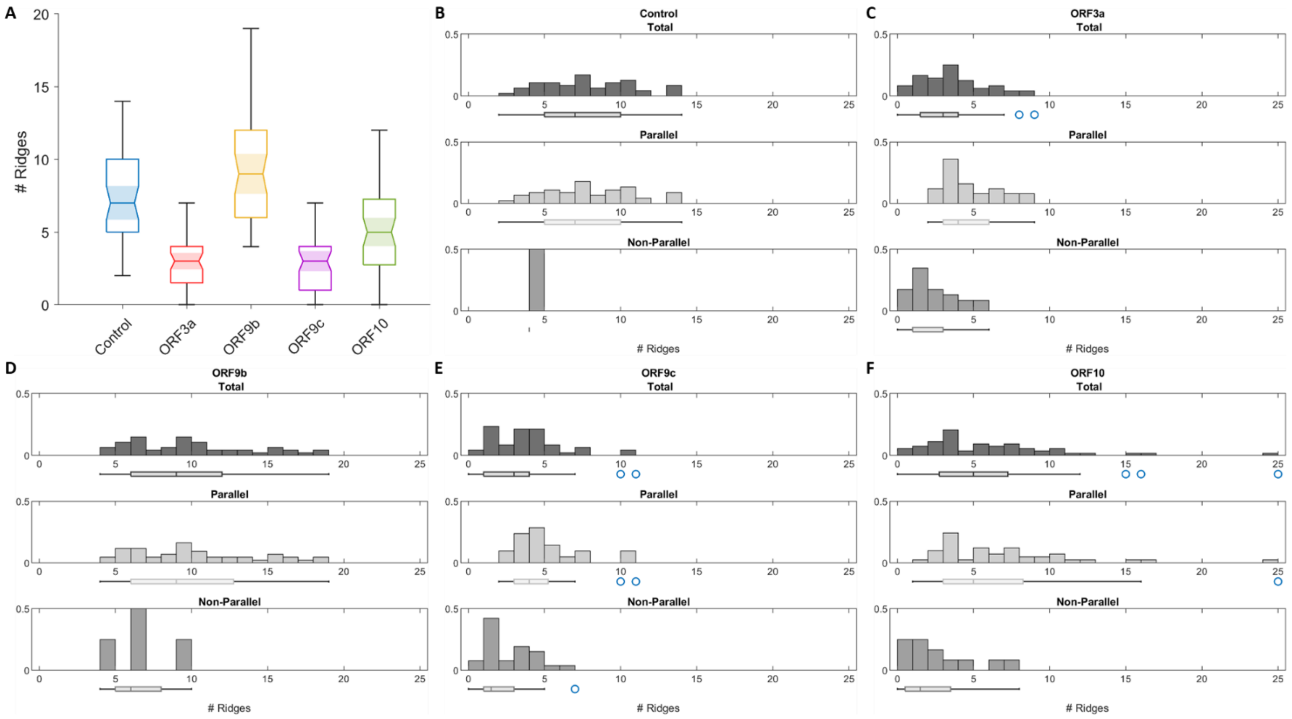


**Figure S4.** Distribution of the number of ridges per mitochondrion evaluated in ORF-A549 cells imaged by transmission electron microscopy. **(A)** Box chart summarizing the distribution of the number of ridges per mitochondrion counted for each cell line. Each box is divided into four sections representing a different quartile while the central lines are the median values and the areas marked in solid colors are the intervals of significance. The full distributions have been depicted in the histograms in **(B-F)**. For each cell line, the corresponding distribution of the number of ridges has been plotted considering the total amount of measurements (upper histograms) as well as separating the mitochondria showing only parallel ridges from the mitochondria with non-parallel ones (middle and lower histograms).

### Supplementary Figure 5

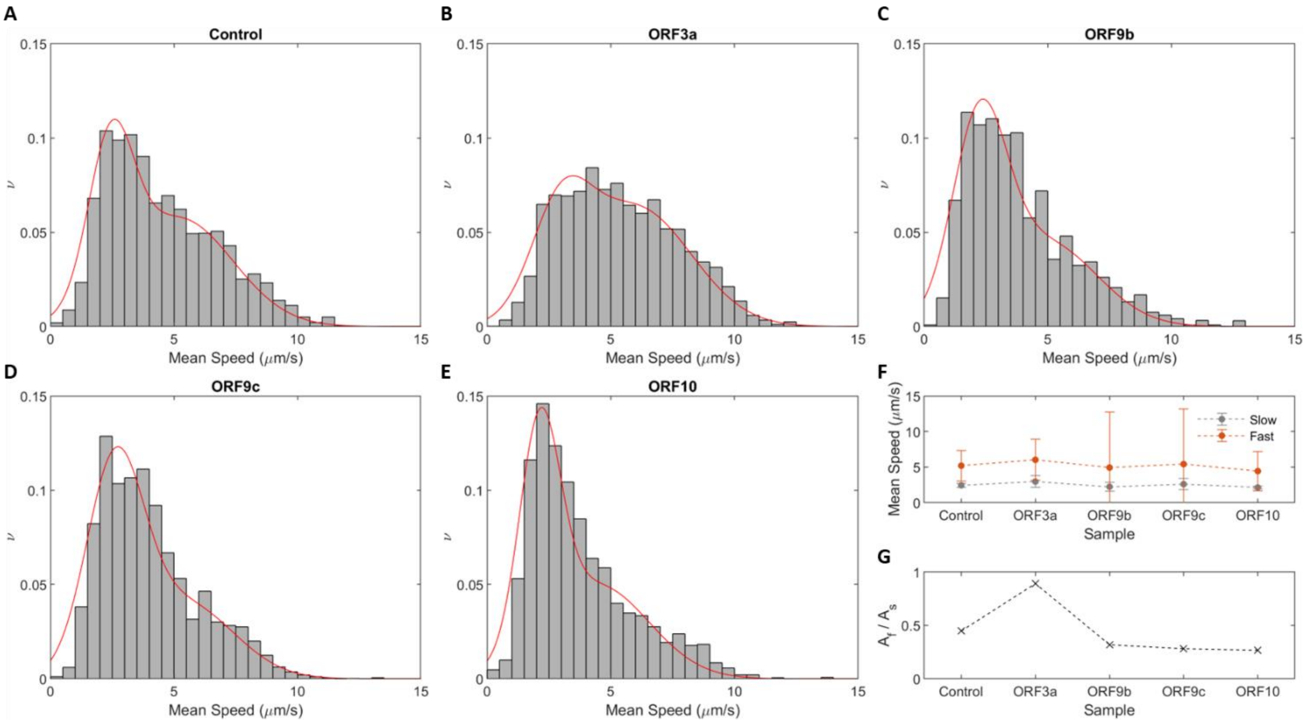


**Figure S5. (A-E)** Distributions of the mitochondria mean speed for each cell line. The red lines in the histograms represent the results of a multiple Gaussian fitting. In all cases, we individuated two Gauss-distributed populations. **(F)** The plot of the fitted mean values of both populations (indicated as “slow” and “fast”) for each cell line. The error bars represent the standard deviations of the Gaussian fitting. **(G)** Ratio between the calculated areas under the Gaussians corresponding to the fast (Af) and slow (As) populations.

#### Supplementary Figure 6

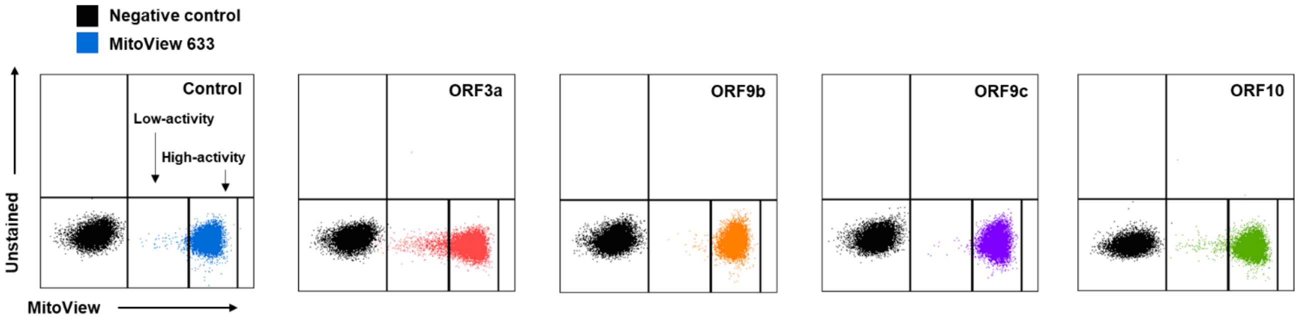


**Figure S6.** Analysis by flow cytometry of mitochondrial activity in ORF-A549 cells. Dotplot of control cells and A549 cells expressing ORF3a, ORF9b, ORF9c or ORF10. Cells were stained with MitoView 633 (5 nM) and showed two mitochondrial populations according to their activity: high-activity and low-activity mitochondria.

#### Supplementary Figure 7

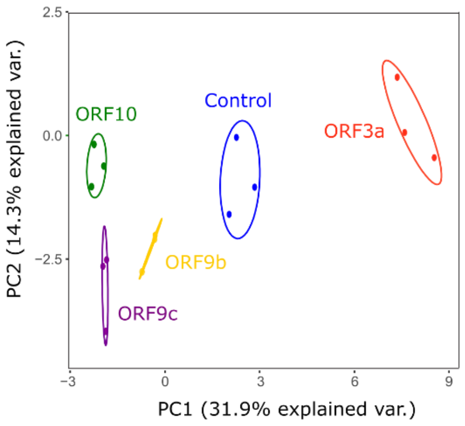


**Figure S7.** Principal Component Analysis (PCA) graph of A549 control cells and A549 cells transduced with ORF3a, ORF9b, ORF9c or ORF10 accessory proteins.

#### Supplementary Figure 8

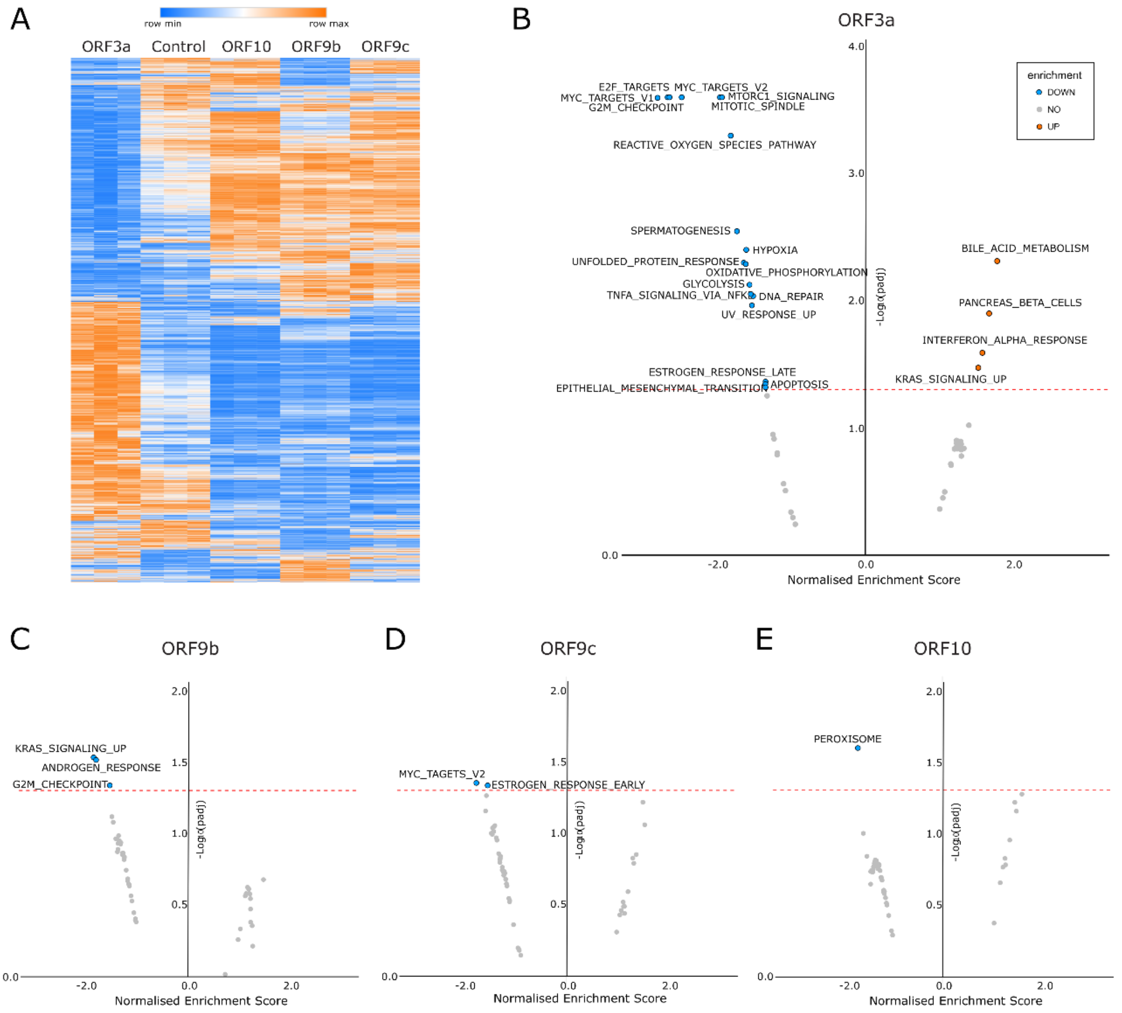


**Figure S8. (A)** Heatmap of RNA-Seq analysis of A549 control cells and A549 cells transduced with ORF3a, ORF9b, ORF9c or ORF10. **(B-E)** Volcano plots of enrichment scores for metabolic pathways, inferred after GeneSet Enrichment Analysis (GSEA) of differentially-expressed genes for ORF3a-A549 **(B)**, ORF9b-A549 **(C)**, ORF9c-A549 **(D)**, and ORF10-A549 cells **(E)**. Horizontal dashed red line represents -log10(padj) = 1.3 (p-value = 0.05).

### Supplementary Figure 9

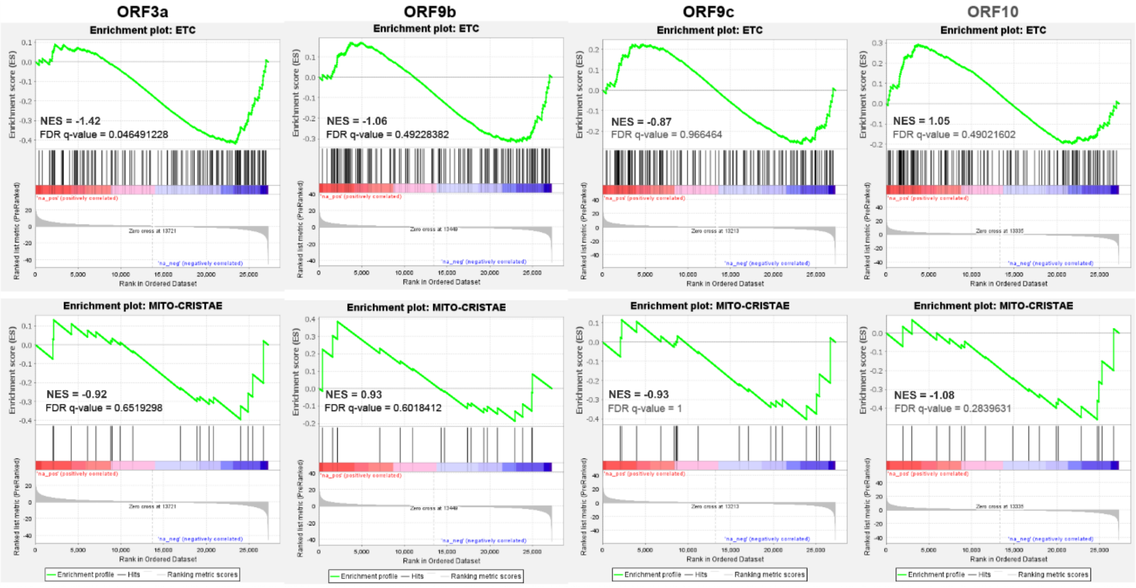


**Figure S9.** Enrichment plots of GeneSet Enrichment Analysis (GSEA) analysis for ETC and mitochondrial cristae gene sets.
